# eIF5A controls mitoprotein import by relieving ribosome stalling at the *TIM50* translocase mRNA

**DOI:** 10.1101/2023.12.19.572290

**Authors:** Marina Barba-Aliaga, Vanessa Bernal, Cynthia Rong, Brian M. Zid, Paula Alepuz

## Abstract

The efficient import of nuclear-encoded proteins into mitochondria is crucial for proper mitochondrial function. The conserved translation factor eIF5A is primarily known as an elongation factor which binds ribosomes to alleviate ribosome stalling at sequences encoding polyprolines or combinations of proline with glycine and charged amino acids. eIF5A is known to impact the mitochondrial function across a variety of species although the precise molecular mechanism underlying this impact remains unclear. We found that depletion of eIF5A in yeast drives reduced translation and levels of TCA cycle and oxidative phosphorylation proteins. We further found that loss of eIF5A leads to the accumulation of mitoprotein precursors in the cytosol as well as to the induction of a mitochondrial import stress response. Here we identify an essential polyproline-containing protein as a direct eIF5A target for translation: the mitochondrial inner membrane protein Tim50, which is the receptor subunit of the TIM23 translocase complex. We show how eIF5A directly controls mitochondrial protein import through the alleviation of ribosome stalling along *TIM50* mRNA at the mitochondrial surface. Removal of the polyprolines from Tim50 rescues the mitochondrial import stress response, as well as the translation of oxidative phosphorylation reporter genes in an eIF5A loss of function. Overall, our findings elucidate how eIF5A impacts the mitochondrial function by reducing ribosome stalling and facilitating protein translation, thereby positively impacting the mitochondrial import process.

---

Mitochondria are complex eukaryotic organelles with endosymbiotic origin (1). In eukaryotes, mitochondria are essential for energy production and macromolecular synthesis as they house key metabolic processes such as oxidative phosphorylation (OXPHOS), electron transport chain (ETC) or TCA cycle, and lipid, amino acids, heme, and Fe-S clusters synthesis. Besides, mitochondria also participate in other cellular processes including Ca^2+^ homeostasis and apoptosis. Given its cellular essentiality, mitochondrial function is critical in health and disease and is a pivotal hallmark of ageing-related disorders (2, 3).

The mitochondrial proteome comprises about 1000 proteins in budding yeast (4). While the mitochondrial genome encodes 1% of them, the other 99% are encoded in the nuclear genome (5). Therefore, these nuclear-encoded mitochondrial proteins, herein termed mitoproteins, need to be transported into the mitochondria to fulfill their biological function. Mitoprotein import is therefore a key process for optimal mitochondrial function (6). Mitoproteins are imported into the mitochondria through both postand cotranslational mechanisms. In the first case, cytosolic chaperones bind to and keep mitoproteins in an unfolded import-competent conformation until delivered to receptors at the mitochondrial surface (7). Co-translational translocation implies the coupling of synthesis and import, with mRNA localization to the mitochondrial surface (8–10).

Protein translocases in the mitochondrial outer and inner membrane (MOM and MIM) mediate the import and sorting of proteins into mitochondria. Mitoproteins initially enter the organelle through the general translocase of the outer membrane (TOM complex), known as the universal entry gate. Most mitoproteins are recognized by their N-terminal positively charged presequences, the most common mitochondrial targeting sequences (MTS) (11). Usually, mitoproteins contact the central receptor Tom20 and cross to the inter-membrane space (IMS) through Tom40, the β-barrel pore-forming subunit. Mitoproteins targeting the MIM or the mitochondrial matrix (MM) are then recognized by the major translocase of the inner membrane (TIM23 complex) with Tim23 as the central subunit and translocase pore and Tim50 as the receptor protein that specifically recognizes the MTS. TIM translocation is energetically driven by the MIM membrane potential and the action of the mtHsp70 motor system. Once reaching their destination, the MTS sequence is cleaved, and the protein is folded. Mitoproteins with no MTS show unclear targeting signals and their import is mediated by other translocases (see reviews (12–14)).

Import failure of mitoproteins leads to proteotoxic effects inside and outside the mitochondria, as unfolded precursors accumulate on the translocases and in the cytosol, which is detrimental to cellular fitness and is associated with various diseases. However, yeast cells are equipped with several stress responses which include a transcriptional and translational reprogramming, reducing the protein synthesis and increasing proteasomal activity to remove accumulated precursors from translocases and cytosol (15–19).

Eukaryotic translation initiation factor 5A (eIF5A) is an essential and highly conserved protein across eukaryotes and archaea (20). eIF5A promotes the translation elongation between amino acids known to be poor substrates for the formation of peptide bonds that may stall translation, such as polyproline motifs but also combinations of proline, glycine, and charged amino acids (21–23). eIF5A is the only known cellular protein containing the post-translational and essential modification hypusine, and in most eukaryotes, it is codified by two highly homologous isoforms, TIF51A and TIF51B in yeast (24, 25). eIF5A expression is regulated according to the metabolic demands of the cell. In yeast, TIF51A gene is up-regulated under respiratory conditions by the transcription (activator/repressor) factor Hap1 (26, 27), which responds to changes in oxygen and heme levels to activate the transcription of many respiratory genes (28). However, under hypoxic/non-respiratory conditions, Hap1 represses TIF51A expression (26, 27).

The cellular adaptation of eIF5A expression according to the energetic status of the cell highlights the essential role of eIF5A in mitochondrial function. Thus, a reduction in eIF5A protein triggers a decrease in the mitochondrial respiration rate and its membrane potential (26, 27, 29–31). In addition, mitochondrial localization of eIF5A has been reported (32–34). Until now, two alternative but related mechanisms to explain the eIF5A role in mitochondrial function have been described. First, eIF5A was reported to control the synthesis and activity of many mitoproteins of macrophage cells. The mitochondrial targeting sequences (MTSs) of some of them were sufficient to confer hypusinated eIF5A-dependent translation efficiency, suggesting that eIF5A might regulate mitochondrial activity by promoting, directly or indirectly, the translation of MTSs of some mitoproteins, which are rich in charged amino acids (35). In connection with this, Zhang et al. (36) reported how eIF5A promotes the translation of some yeast respiratory proteins in response to oxygen levels and suggested that eIF5A and its hypusine would favor a suitable interaction between the amino acids from the MTS regions and the peptide exit tunnel. Here, we describe an alternative molecular mechanism by which eIF5A promotes mitochondrial activity. This mechanism relies on the eIF5A-dependent translation of Tim50, the receptor subunit of the TIM23 complex, impacting the mitochondrial import process. Through the alleviation of ribosome stalling of TIM50 mRNA and thus, modulation of Tim50 protein levels, eIF5A specifically controls the import and translocation of mitochondrial proteins. Hence, the loss of eIF5A drives the accumulation of precursors in the cytosol, activates the mitochondrial import stress response while secondarily reduces the translation and levels of many mitochondrial proteins.

## Results

### Depletion of eIF5A results in the down-regulation of many mitochondrial proteins including components of OXPHOS, TCA and mitochondrial membrane transporters

We and others have found that eIF5A is necessary to maintain high mitochondrial activity; additionally, several mitoproteins whose levels drop in the absence of functional eIF5A have been identified in yeast and mammalian cells (26, 27, 35, 36). In order to expand our knowledge on the role of eIF5A in mitochondria, we performed a proteomic experiment in which a wild-type and two eIF5A temperature-sensitive mutant yeast strains, *tif51A-1* carrying a single point mutation (Pro83 to Ser) and *tif51A-3* carrying a double point mutation (Cys39 to Tyr, Gly118 to Asp) (37, 38), were exponentially grown in YPD media at 25°C and then incubated at 41°C for 4 h to deplete eIF5A. Data from the three replicates shown in the MDS-plot indicated that eIF5A mutants are already different in their proteome with respect to the wild-type at permissive temperature, and this difference is exacerbated at restrictive temperature (Fig EV1A). We also confirmed the response of wild-type to heat stress conferred by incubation at 41°C as there was an enrichment in the functional categories (GOs) of ‘Cellular response to stress’ in its up-regulated proteins (Fig EV1B). We calculated the 41°C/25°C fold change of protein levels and then, the ratios between this fold change for each mutant respect to the wild-type. Out of the 1358 proteins detected in the three strains, 292 were significantly down-regulated and 135 up-regulated in at least one mutant respect to the wild-type (138 down- and 38 up-regulated in both eIF5A mutants). Down-regulated proteins were enriched in several functional categories related to mitochondria (Fig EV1C and Table S1), including OXPHOS proteins and TCA enzymes (Figs 1A,B and Table S1). We observed that several mitochondrial membrane transporters were also down-regulated, as the phosphate (*Mir1*) and the ADP/ATP (*Pet9*) MIM carrier proteins and members of the TOM translocase of the MOM (*Tom20, Tom70*) (Figs 1A,B, EV1C and Table S1). To independently corroborate the proteomic results, we confirmed the reduction in the levels of two of the most down-regulated mitoproteins, the *Por1* porin of the MOM and the *Hsp60* chaperone, in both *tif51A-1* and *tif51A-3* mutants at restrictive temperature (Figs 1C-E). On the other hand, proteins of functional categories related to cytoplasmic translation were up-regulated in eIF5A mutants with respect to wild-type due to a lower decrease in the 41°C/25°C fold change in the eIF5A mutants (Figs 1A,B and EV1C). Proteins of the category ‘drug membrane transport’ (*Snq2, Yor1, Qdr2, Pdr5*) were also up-regulated in the eIF5A mutants at 41°C (Figs 1A, EV1C and Table S1). In sum, the proteomic results indicated, among other effects, the requirement of eIF5A to maintain adequate protein levels of a wide range of mitoproteins.

**Fig. 1.**
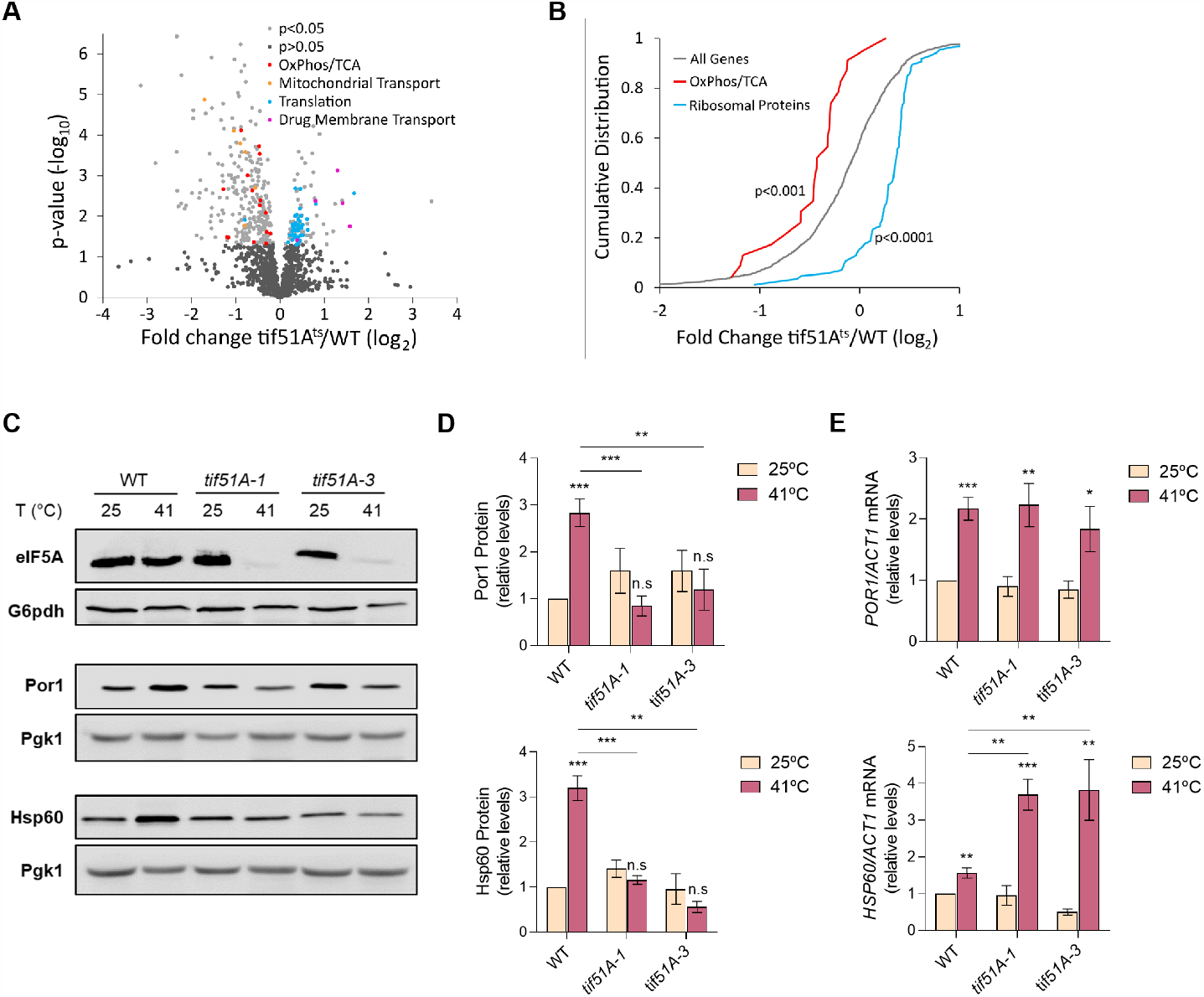
Proteomic analysis of eIF5A deletion shows down-regulation of mitochondrial proteins. (A) Volcano plot showing the log2 fold change values of 1358 detected proteins plotted against their associated log10 p-values. Dots representing individual proteins were divided in five different groups: p-value > 0.05 (black); p-value < 0.05 (grey); OXPHOS/TCA proteins (red); mitochondrial transport proteins (orange); translation proteins (blue) and drug membrane transport proteins (magenta). (B) Cumulative distributions of log2 fold changes of all proteins detected (grey), OXPHOS/TCA (red) and ribosomal proteins (blue). (C,D) Wild-type, *tif51A-1* and *tif51A-3* strains were cultured in YPD at 25°C until early exponential phase and transferred to 25°C or 41°C for 4 h. eIF5A, Por1 and Hsp60 protein levels were determined by western blotting (C) and quantified (D). G6PDH levels were used as loading control. A representative image is shown. (E) *POR1* and *HSP60* mRNA relative levels were determined by RT-qPCR. Data information: In (D,E) Results are presented as mean ± SD from three independent experiments. The statistical significance was measured by using a two-tailed paired Student t-test relative to 25°C. *p < 0.05, **p < 0.01, ***p < 0.001.

An important outcome of our proteomic analysis was that, among the down-regulated mitoproteins upon eIF5A depletion, most of them do not contain consecutive prolines nor a high number of other eIF5A-dependent motifs in their sequences (22, 23) (Table S1). This finding had been previously observed in yeast and mammalian cells (35, 36) and suggests a general down-regulation of mitochondrial proteins levels in the absence of eIF5A and, thus, a role for eIF5A different to that well-described and direct one on translating problematic amino acids for peptide bond formation.

### eIF5A depletion causes a down-regulation of mitochondrial proteins translation

The specific decrease in the levels of mitoproteins upon eIF5A depletion could be explained by decreased translation of the corresponding mRNAs, as it is suggested by similar or even higher mRNA levels of the down-regulated proteins Por1 and Hsp60 in *tif51A-1* and *tif51A-3* mutants (Fig 1E). To further investigate if the general down-regulation of mitoproteins was occurring at the translational level, we obtained polysome profiles for wild-type and *tif51A-1* strains in galactose media to promote respiration at both permissive (25°C) and restrictive (37°C) temperatures and determined the distribution of specific mRNAs among the different polysomal fractions. Importantly, polysomes were maintained in both strains at both temperatures, indicating that global translation was not significantly affected (Figs 2A,B). To analyse the distribution of short length-RNAs along the fractions, we divided the polysome profiles in three different sections corresponding to the monosomal fractions (M), fractions occupied by mRNAs with 2 or 3 ribosomes (2n-3n) and the rest of the polysomal fractions (P). For long length-RNAs we used the sections corresponding to the monosomal fractions (M), fractions with 2 to 8 ribosomes (2n-8n) and the rest of the polysomal fractions (P) (Fig EV2A). In agreement with unaffected global translation, we found no differences in the ratios P/2n-3n and P/2n-8n between the two strains at both permissive and restrictive temperatures (Figs EV2B,C).

**Fig. 2.**
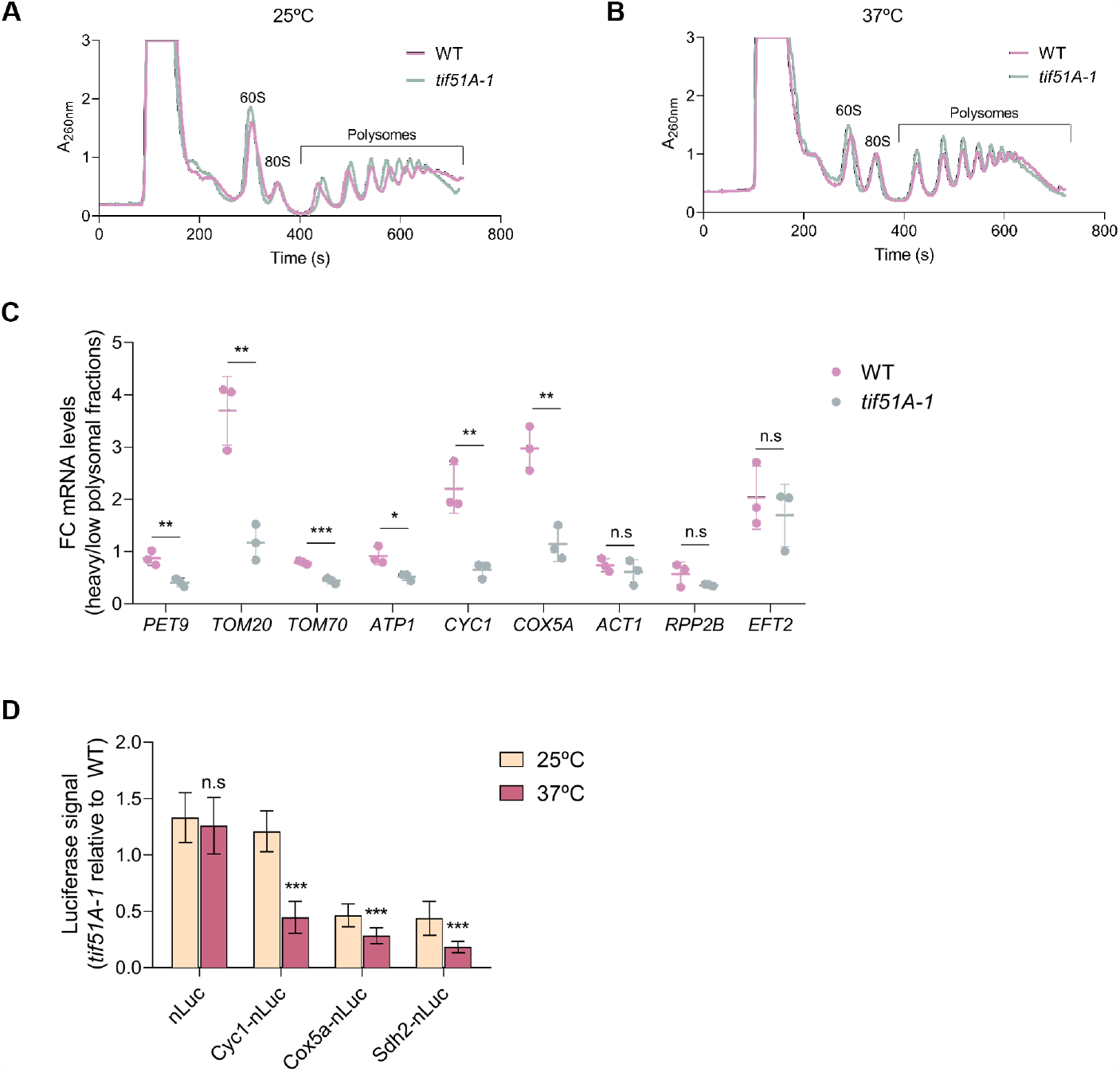
The levels of mitochondrial proteins are post-transcriptionally affected upon eIF5A depletion. (A,B) Polysome profiles were obtained for wild-type and *tif51A-1* strains cultured in SGal at 25°C (A) or 37°C (B) for 4 h. (C) The RNA from individual fractions of the polysomes profiles was extracted and the mRNA levels were analyzed by RT-qPCR using gene specific primers in the corresponding sections at restrictive temperature. Fold change between the heavy and the low polysomal sections are presented. (D) Wild-type strain and *tif51A-1* expressing nLuc, Cyc1-nLuc, Cox5a-nLuc or Sdh2-nLuc (from left to right) reporters were cultured in YPD until early exponential phase and then transferred to 25°C or 37°C for 4 h. After addition of doxycycline to induce luciferase expression, the luminescence levels generated by the nanoluciferase 60 mins after the addition of the furimazine substrate were measured. Data information: In (C,D) Results are presented as average ± SD from three independent experiments. The statistical significance was measured by Student t-test relative to wild-type strain or to 25°C. *p < 0.05, **p < 0.001, ***p < 0.001.

Next, we investigated the distribution of mRNAs encoding for mitoproteins with different biological function and intra-mitochondrial localization. We calculated the fold change between the mRNA abundance found at heavy and low polysomal fractions. Significant lower ratios were observed for *PET9, TOM20* and *TOM70* mRNAs encoding mitochondrial transporters at the MIM and MOM in the *tif51A-1* strain at restrictive temperature (Figs 2C and EV2D) with no significant differences at permissive temperature (Fig EV2G). These results indicated that the mRNA abundance was shifted to earlier fractions and translation was reduced. Similar results were observed for the *ATP1, CYC1* and *COX5A* mRNAs encoding OXPHOS components at the MIM and IMS (Figs 2C and EV2E,H). The reported lower ribosome association with the mitochondrial mRNAs upon eIF5A depletion was in agreement with our proteomic analyses, since 7 out of the 8 tested mRNAs encoding for mitoproteins were detected in the proteomic study and showed down-regulation of the protein levels in the two eIF5A mutants at restrictive temperature (Table S1).

We also studied the translation of the constitutively expressed *ACT1, RPP2B* and *EFT2* mRNAs encoding actin, a ribosomal protein and translation elongation factor, respectively, localized in the cytosol. Similar ratios were found between the wild-type and *tif51A-1* strains at both temperatures, indicating that the translation of these mRNAs remained unaffected in the absence of eIF5A (Figs 2C and EV2F,I), which agrees with the finding that global translation is not affected in the *tif51A-1* mutant (Figs EV2B,C).

To specifically address whether the synthesis of mitoproteins by translation is down-regulated upon eIF5A depletion, we used a reporter construct integrated in the *URA3* locus, consisting of a *tetO7* inducible promoter and a nanoluciferase (nLuc) reporter ORF for measuring kinetics of protein synthesis. This system was recently used to quantify protein levels and translation stall duration from reporter mRNAs in yeast (39). We generated fusions of the OXPHOS genes *CYC1* and *COX5A*, and the TCA enzyme *SDH2* to the nLuc reporter gene. After inducing expression, we analysed the protein synthesis by incubating for 1 h in anhydrotetracycline-supplemented media. We did not find significant differences in protein synthesis between the *tif51A-1* mutant and the wild-type strain at restrictive temperature for the nLuc reporter alone, which targets the cytosol (Fig 2D). However, the synthesis of Cyc1, Cox5a and Sdh2 luciferase fusions was significantly affected at restrictive temperature in the *tif51A-1* strain (Fig 2D). The obtained results point towards a specific mechanism connected to eIF5A to coordinately reduce the synthesis of mitoproteins at the translation level and, therefore, mitochondrial protein levels. However, this mechanism seems to not be connected to the presence of peptides requiring eIF5A for their synthesis during translation as none of the investigated mitochondrial mRNAs encode for polyprolines nor a high number of other eIF5A-dependent motifs.

### eIF5A depletion induces the mitochondrial protein import stress mitoCPR and MitoStores

The “drug membrane transport” functional group, which includes the ATP-binding cassette (ABC) transporters Pdr5, Snq2 and Yor1, was found among the few GOs up-regulated in the eIF5A mutant in our proteomic study (Figs 1A, EV1C and DATA). The evolutionarily conserved transporter *PDR5* has been shown to be induced during the mitochondrial stress response called mitoCPR (mitochondrial compromised protein import response) (18). mitoCPR is induced upon defects in the translocation and import of mitoproteins to increase the activity of the proteasome in the cytosol, remove accumulated precursors from the TOM complex and stabilize homeostasis. This response relies on the Pdr3-mediated transcriptional activation of different genes related to the multidrug resistance (MDR) response, such as *PDR5*, which is involved in lipid metabolism and transport. The Pdr3-induced expression of Cis1, among others, recruits the AAA-protease Msp1 to the TOM translocase of the MOM to mediate the removal of accumulated mitoproteins upon import failure (18). To investigate if the mitoCPR response was induced upon eIF5A depletion, we analysed the mRNA expression of *PDR3* and the Pdr3-responsive genes *MSP1, GRE2, PDR1, PDR3, PDR5* and *CIS1* in the *tif51A-1* mutant at restrictive temperature. The *tif51A-1* mutant, but not the wild-type, showed a significant up-regulation of all genes except *MSP1*, with *CIS1* and *PDR5* the most induced mRNAs (Fig 3A).

**Fig. 3.**
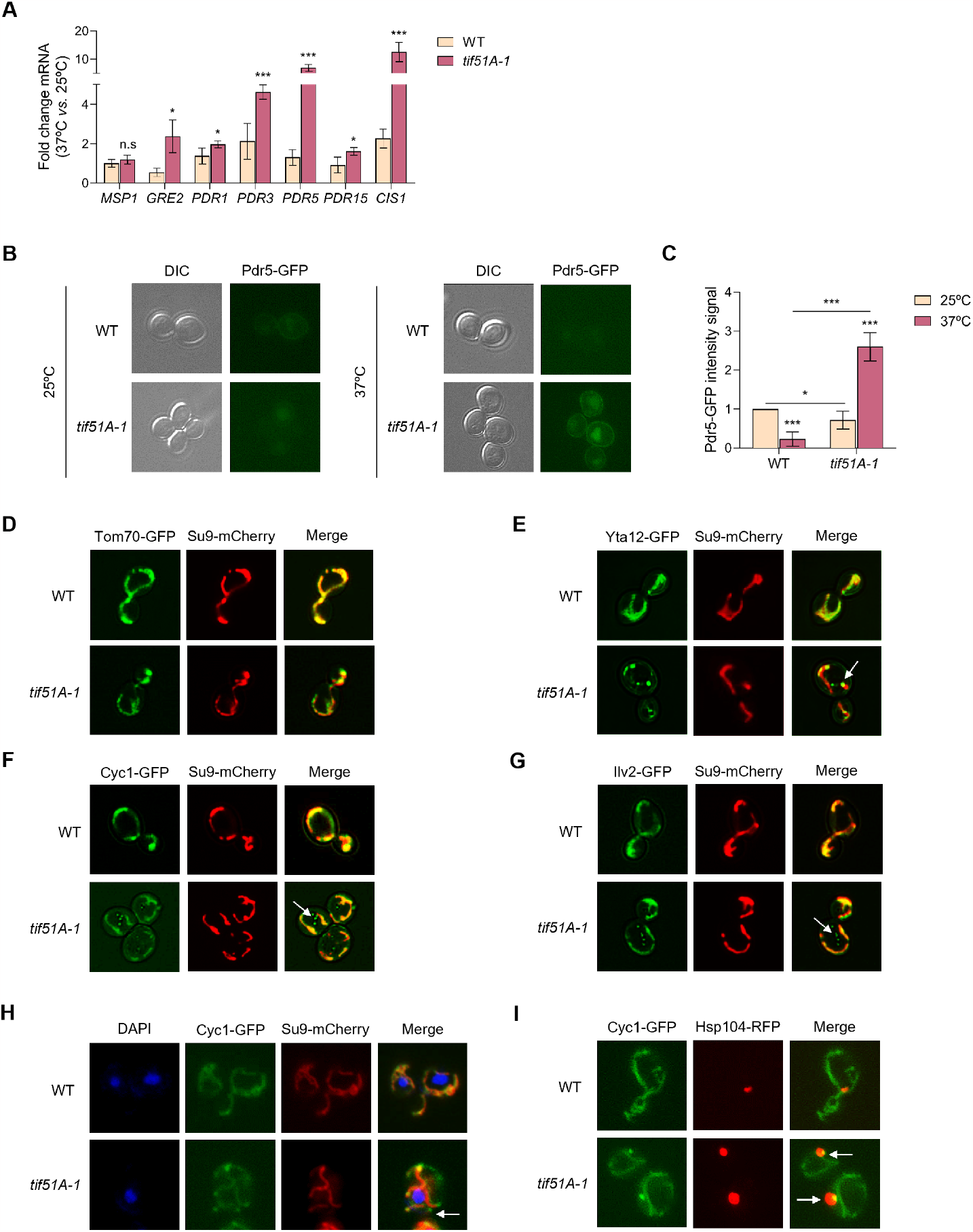
eIF5A deficiency generates mitoCPR stress and mislocalization of mitoproteins. (A) Wild-type strain and *tif51A-1* were cultured in SGal medium at 25°C until reaching post-diauxic phase and then transferred to 25°C or 37°C for 4 h. mRNA relative levels from mitoCPR genes were determined by RT-qPCR. (B) Wild-type strain and *tif51A-1* expressing Pdr5-GFP were cultured as in (A) and then subjected to fluorescence microscopy. (C) Quantification of Pdr5-GFP fluorescent signal from at least 150 cells. (D-G) Wild-type strain and *tif51A-1* expressing Tom70-GFP (D), Yta12-GFP (E), Cyc1-GFP (F) or Ilv2-GFP (G) and Su9-mCherry were cultured as in (A) and subjected to fluorescence microscopy. (H) Wild-type strain and *tif51A-1* expressing Cyc1-GFP and Su9-mCherry were cultured as in (A) and incubated for 30 mins with DAPI prior microscopy to stain the nuclei. (I) Wild-type strain and *tif51A-1* expressing Cyc1-GFP and Hsp104-RFP were cultured as in (A). (B,D-I) A representative image is shown from three independent experiments. Data information: In (A,C) Results are presented as mean ± SD from three independent experiments. The statistical significance was measured by using a two-tailed paired Student t-test relative to 25°C. *p < 0.05, ***p < 0.001.

When induced by Pdr3, the Pdr5 ABC membrane transporter is observed in the plasmatic membrane, where it mediates the efflux of xenobiotics, as well as in the vacuole, where it is degraded (**?**). We fused the Pdr5 ORF to GFP and analysed its cellular localization by fluorescence microscopy. We found that the levels of Pdr5 were almost non-detectable in the wild-type strain at both temperatures as well as in the *tif51A-1* strain at permissive temperature. However, at restrictive temperature, Pdr5 protein levels were significantly induced in the *tif51A-1* strain and could be observed at their expected cellular locations (Figs 3B,C). Then, we added Nile Red, a Pdr5 specific substrate which stains lipid granules, to the cell cultures. We observed red fluorescent signal inside the cells in all the tested conditions except for the *tif51A-1* mutant at restrictive temperature, indicating that high Pdr5 pumping activity excludes the Nile Red from these cells (Figs EV3A,B).

Along with a transcriptional response, it has been found that mitoproteins can aggregate in the cytosol upon induction of mitochondrial import stress (40, 41). To explore the potential mislocalization of mitoproteins targeted to different locations within the mitochondria following eIF5A depletion, we investigated their distribution relative to a Su9-mCherry marker, which served to visualize the overall mitochondrial network. Tom70 is part of the TOM complex, placed in the MOM. Cytochrome c isoform 1 (Cyc1) is involved in the transfer of electrons during cellular respiration and targets the IMS. Yta12 is an m-AAA protease component required for the degradation of misfolded or unassembled proteins and targets the MIM. Ilv2 is an acetolactate synthase involved in the synthesis of isoleucine and valine and targets the MM. We found that the outer membrane protein Tom70 shows no difference between the wild-type and *tif51A-1* mutant at permissive and restrictive temperatures (Figs 3D and EV3C). Figs 3E-G show that at restrictive temperature Yta12, Cyc1 and Ilv2 are localized in the mitochondria in the wild-type strain but, in the *tif51A-1* mutant they also form foci, which do not co-localize to the mitochondria. On the contrary, no foci were observed at permissive temperature (Figs EV3D-F). Although Ilv2 has been described to accumulate as one big aggregate in the nucleus upon protein import failure (42), we found that in the absence of eIF5A, this protein accumulates in multiple and distributed foci, similarly to the misslocalized protein aggregates of Cyc1 and Yta12.

Next, we asked about the intracellular localization and composition of these eIF5A-dependent mitoprotein foci. Different cellular destinations for mitoproteins aggregates have been described including the cytosol, endoplasmic reticulum and nucleus (42). DAPI staining of the nucleus of the cells indicated that Cyc1 aggregates were not co-localizing in the nucleus and thus, seemed to be part of the cytosol (Fig 3H). To gain insight into the accumulation of non-imported mitochondrial precursors in cytosolic granules upon eIF5A depletion, we investigated whether chaperones such Hsp104 could be controlling this process. Hsp104 is a disaggregase that binds to aggregated or misfolded proteins and disentangles them in an ATP-dependent manner (43–45). Recently it was found that induction of mitochondrial import stress can cause cytosolic accumulation and co-localization of mitochondrial matrix-destined precursor proteins with Hsp104 granules. We fused Hsp104 ORF to RFP in a strain already expressing Cyc1-GFP and analysed its cellular localization. We found that a high percentage (>80%) of Hsp104 foci colocalize with Cyc1 aggregates (Fig 3I).

Altogether, these results suggest that depletion of eIF5A compromises mitochondrial protein import, as the induction of the Pdr3-mediated mitoCPR response and the cytosolic Hsp104 aggregation of precursor mitoproteins into Mito-Stores have been described as mechanisms to reduce toxicity from accumulated mitoprotein precursors and to restore cellular homeostasis upon import failure.

### eIF5A alleviates ribosome stalling of TIM50 mRNA encoding the mitochondrial inner membrane receptor

Given the elevated mitochondrial protein import stress response found upon eIF5A depletion, we sought to explore the mechanism by which eIF5A impacts the mitochondrial protein import. Tim50 protein is a receptor component of the MIM translocase complex TIM23 which recognizes the N-terminal MTS-containing proteins after emerging from the TOM complex and mediates their import to the mitochondrial inner membrane and matrix ((46), see review (47)). Interestingly, Tim50 contains a 10 proline residues in a 14 amino acid region in the first half of the protein with 7 of them consecutive (Fig 4A) and, therefore, is a proposed putative eIF5A-target (27). In addition, *TIM50* overexpression has previously been found to induce mitochondrial protein import defects (18). We therefore sought to investigate the possible role of eIF5A in *TIM50* mRNA translation. First, we genomically attached an HA tag to Tim50 to quantify the impact of eIF5A depletion on endogenous Tim50 expression. We found a significant drop in Tim50 protein levels upon eIF5A depletion with no decrease in its mRNA levels (Fig EV4A). Moreover, expressing a *FLAG-TIM50-GFP* version under a *GAL* inducible system, we detected a reduction in Tim50 protein synthesis in the two eIF5A mutants, that was not due to lower levels of the corresponding mRNA, further suggesting a role of eIF5A in the translation of *TIM50* mRNA (Fig EV4B).

**Fig. 4.**
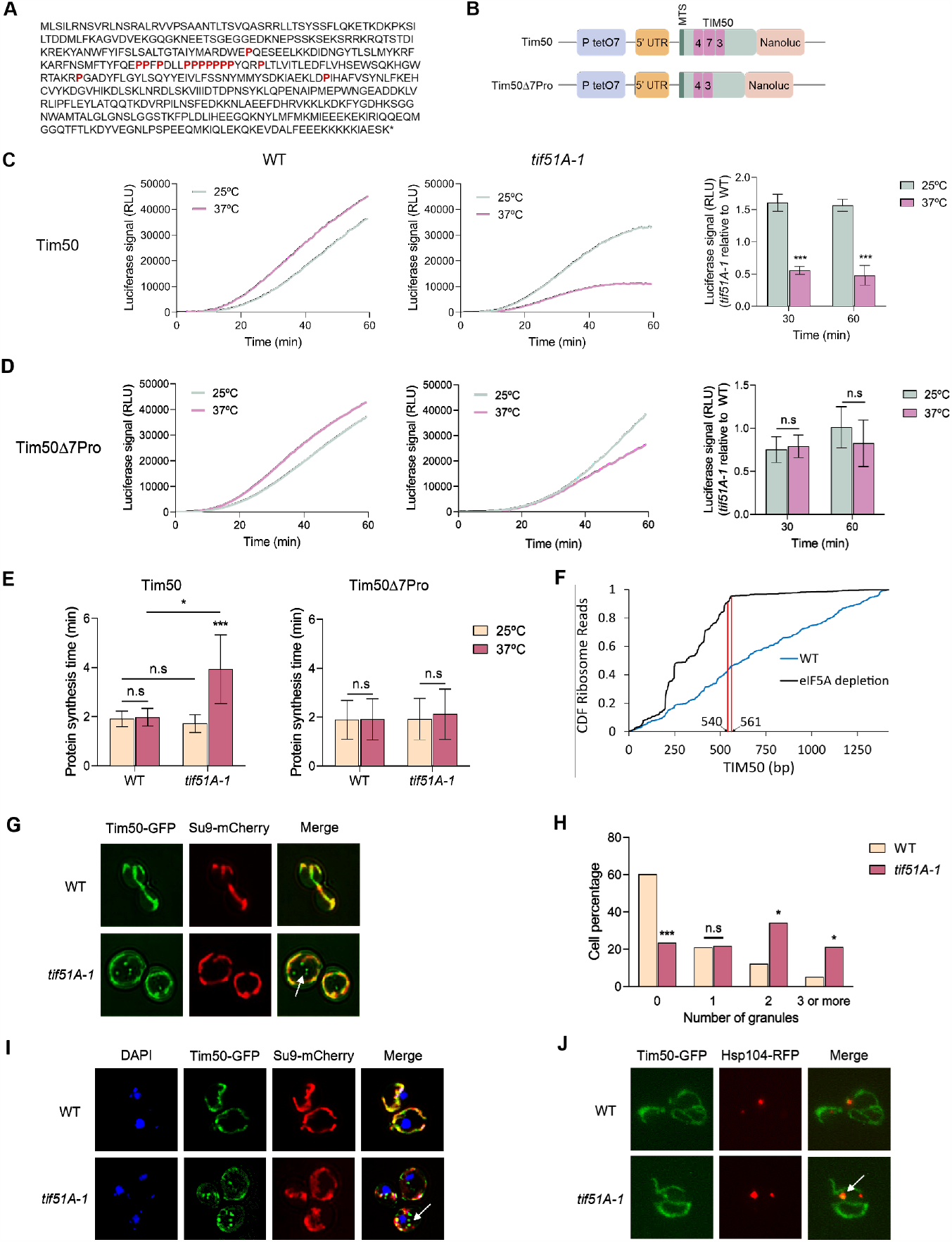
eIF5A depletion causes TIM50 polyproline ribosome stalling, decreased Tim50 protein levels and mislocalization. (A) Scheme showing Tim50 protein sequence. (B) Scheme showing the two different nLuc versions used in this study. Proline numbers are shown in magenta. (C,D) Wild-type strain (left) and *tif51A-1* (middle) expressing the wild-type Tim50 (B) or Tim507Pro (C) were cultured in YPD until early exponential phase and then transferred to 25°C or 37°C for 4 h. After addition of doxycycline to induce luciferase expression, the luminescence levels generated by the nLuc after the addition of the furimazine substrate were measured along time. A representative experiment is shown. Quantification of the Tim50 protein levels is shown in the right. (E) Full protein synthesis time was calculated for wild-type strain and *tif51A-1* expressing the wild-type Tim50 (left) or Tim507Pro (right). (F) Fraction of ribosome reads of various lengths along TIM50 transcript in (23) ribosome profiling libraries. (G) Wild-type strain and *tif51A-1* expressing Tim50-GFP and Su9-mCherry were cultured in SGal medium at 25°C until reaching post-diauxic phase, transferred to 37°C for 4 h and subjected to fluorescence microscopy. (H) Quantification of cells with Tim50 aggregates at 37°C is shown. (I) Wild-type strain and *tif51A-1* expressing Tim50-GFP and Su9-mCherry were cultured as in (G) and incubated for 30 mins with DAPI prior microscopy to stain the nuclei. (J) Wild-type strain and *tif51A-1* expressing Tim50-GFP and Hsp104-RFP were cultured as in (G). (G-I,J) A representative image is shown from three independent experiments. Data information: In (C-E,H) Results are presented as mean ± SD from three independent experiments. Statistical significance was measured using a two-tailed paired Student t-test relative to 25°C. *p < 0.05, ***p < 0.001.

Tim50 protein half-life is approximately 9.6 h (48), which makes it more difficult to measure large differences in new protein synthesis. Therefore, to accurately test and quantify the eIF5A-dependency for *TIM50* mRNA translation, we fused two different versions of the *TIM50* DNA sequence to a TetO7-inducible nLuc reporter: the first one expressing the wild-type *TIM50* sequence and the second one expressing *TIM50* with a deletion of the 7 consecutive prolines (Tim507Pro) (Fig 4B). After inducing expression, we analysed the protein synthesis by incubating for 1 h in anhydrotetracycline-supplemented media the wild-type and mutant cells. After analysing the protein synthesis, it was observed that the *tif51A-1* mutant showed a 3-fold reduction in the synthesis of Tim50 at restrictive temperature compared to the wild-type strain, further suggesting Tim50 as an eIF5A target for translation elongation (Fig 4C). Considering that final protein abundance is determined by both mRNA levels and translation, we confirmed that the lower protein levels were not attributed to *TIM50* transcription, since the mRNA levels were even slightly higher in the *tif51A-1* mutant at restrictive temperature compared to wild-type (Fig EV5A). To test if the decrease in protein synthesis was due to the presence of a high number of prolines in the Tim50 sequence, we analysed the Tim50 version containing deletions of the 7 consecutive prolines. Results indicated that after removing the proline-rich region of Tim50, protein synthesis is rescued in the *tif51A-1* mutant at 37°C and reaches similar protein levels to the wild-type strain (Fig 4D).

Inducible reporter assays enabled us to calculate both the overall expression levels as well as the time needed for translation elongation through the region upstream of the nLuc reporter gene (39, 49). By using the Schleif plotting technique (50, 51), we compared the amount of time needed to detect luciferase signal from a reporter only expressing nLuc and the two Tim50 reporters. Then, we obtained the time required for ribosomes to translate each of the Tim50 versions. We found that the time required for translating a wild-type *TIM50* mRNA was significantly increased in the *tif51A-1* mutant at restrictive temperature. If a cell needs approximately 2 minutes to generate one full Tim50 polypeptide, upon eIF5A depletion, the ribosomes need almost 4 minutes to achieve the complete translation of this mRNA. However, the synthesis time required for translating TIM507Pro mRNAs were almost identical to those in the corresponding wild-type strain (Fig 4E). This extended translation duration was indicative of a ribosome stall at the polyprolines. To further confirm ribosome stalling, we analysed the ribosome density across the TIM50 transcript in control and an eIF5A degron strain (23). We found a precipitous drop-off in ribosome density exactly where the stretch of polyprolines is located in Tim50 (540-561bp) upon eIF5A depletion but not in the control strain (Fig 4F). This absence of ribosomes after the stretch of polyprolines is indicative of ribosome stalling and the induction of ribosome quality control at the polyproline residues.

*TIM50* mRNA is constitutively localized to the vicinity of the mitochondrial surface and the import of Tim50 protein to mitochondria occurs co-translationally (9, 10). To confirm that ribosome stalling along *TIM50* mRNA upon eIF5A depletion was mitochondrially localized, we visualized mitochondria using the marker Su9-mCherry and *TIM50* mRNAs with the MS2 tag system (10) in both wild-type and *tif51A-1* strains at restrictive temperature. We observed *TIM50* mRNA to be strongly associated with the mitochondrial surface in both strains (Fig EV5B). This result indicates that eIF5A depletion does not affect the localization of *TIM50* mRNA molecules and that Tim50 ribosome stalling occurs adjacent to the mitochondrial surface.

Finally, we asked if ribosome stalling along *TIM50* mRNA upon eIF5A depletion could affect its protein localization. We observed that Tim50 strongly localizes to the mitochondria in the wild-type strain. However, we found the presence of Tim50 foci which do not co-localize to the mitochondria in the *tif51A-1* mutant whereas no aggregates were observed at permissive temperature (Figs 4G and EV5C). The number of cells containing more than one of these Tim50 aggregates was found to be significantly increased in the *tif51A-1* mutant at restrictive temperature (Fig 4H). DAPI staining of the nucleus of the cells indicated that Tim50 aggregates were not co-localizing in the nucleus and thus, seemed to be part of the cytosol (Fig 4I). To gain insight into the accumulation of Tim50 into Mitostores, as previously seen for Cyc1, we fused Hsp104 ORF to RFP in a strain already expressing Tim50-GFP and found that a high percentage (>70%) of Hsp104 foci co-localize with Tim50 aggregates (Figs 4J and EV5D). Together, these results indicated that Tim50 down-regulation in the *tif51A-1* mutant was due to a decrease in translation because ribosomes stall along the proline-rich region of its sequence. Thus, these data strongly suggest that *TIM50* mRNA translation is directly dependent on eIF5A and its deficiency generates ribosome stalling on the mitochondrial surface and precursor protein accumulation in the cytosol.

### Deletion of Tim50 proline stretch in eIF5A mutant restores mitochondrial protein import without rescuing mitochondrial respiration

Herein, we have demonstrated how upon eIF5A depletion, Tim50 translation occurring at the mitochondrial surface stalls in the polyproline stretch and its protein synthesis is decreased. This generates specific import defects for mitoproteins, which accumulate and aggregate in the cytosol. Therefore, we asked if a reduction in Tim50 stalling and rescuing of Tim50 protein levels would alleviate the import collapse and its derived effects associated with eIF5A loss of function. To test this, we generated two strains expressing the endogenous Tim50 protein without the region containing the 7 consecutive prolines of its sequence (Tim50Pro) fused to GFP, the first one in a wild-type (BY4741) background and the second one in a *tif51A-1* mutant background. In these strains, the endogenous *TIM50* gene is mutated so that the only source of Tim50 protein for the cell is the eIF5A-independent version Tim50Pro. Here, translational stalling of *TIM50* mRNA occurring at the mitochondrial surface is no longer present and Tim50 reporter levels are rescued upon eIF5A depletion (Figs 4C-E). Surprisingly, the functionality of the Tim50Pro version was shown to be almost unaffected. We observed no obvious growth differences in glycerol media between the wild-type strain containing full Tim50 or Tim50Pro (Fig 5A), although the proline-rich region is found in the presequence-binding groove of the Tim50 IMS domain, which functions in receiving the proteins in the TIM23 complex (52). Therefore, the proline-rich region is not essential for cell viability under the respiratory conditions tested and thus, for the proper mitochondrial import.

**Fig. 5.**
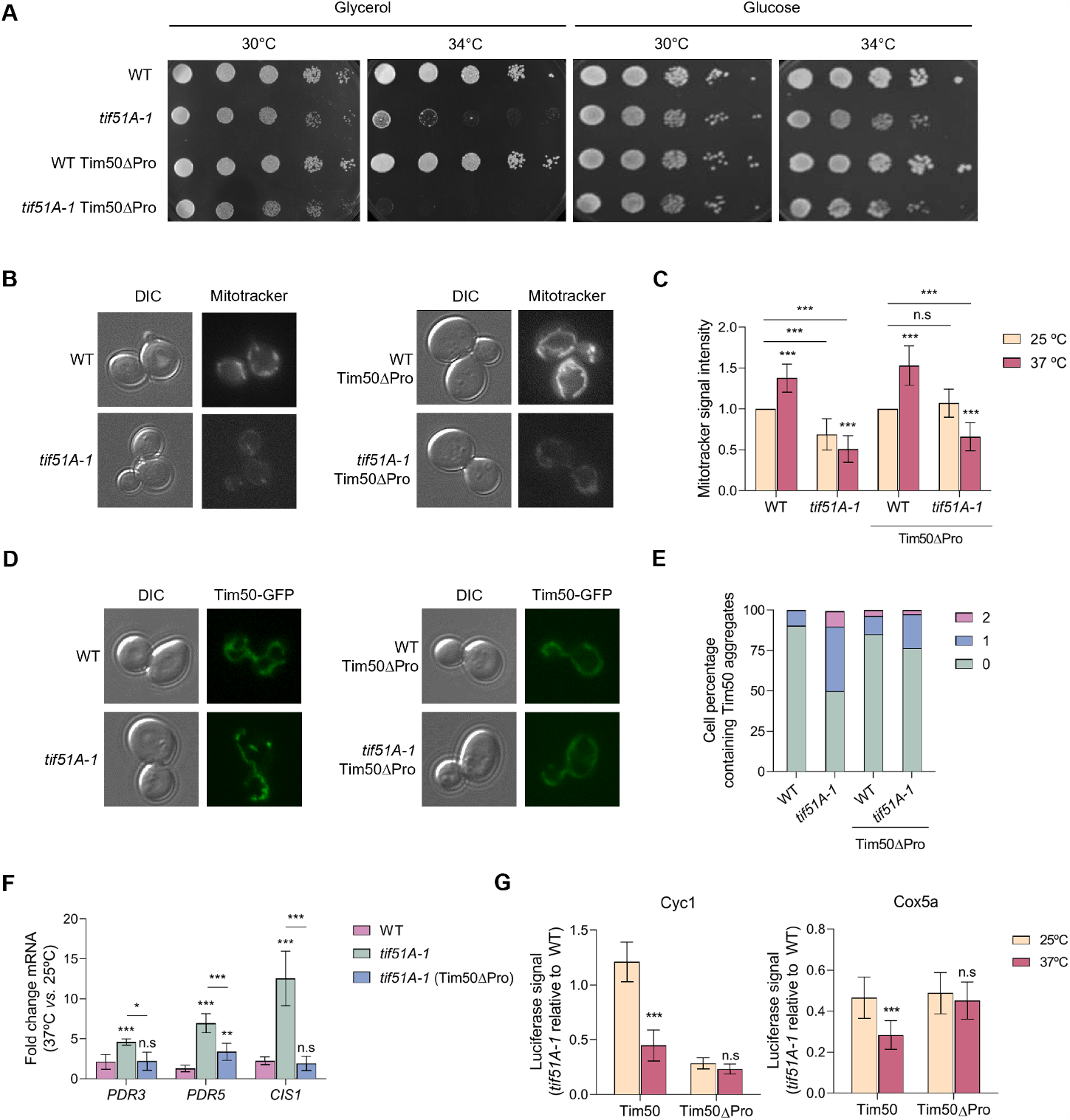
Deletion of the Tim50 polyproline stretch in the eIF5A mutant does not rescue mitochondrial respiration but cancels the mitoCPR response induction. (A) Growth of the wild-type, *tif51A-1*, wild-type Tim50Pro and *tif51A-1* Tim50Pro was tested in YPGly and YPD media at the indicated temperatures. (B-E) Wild-type, *tif51A-1*, wild-type Tim50Pro and *tif51A-1* Tim50Pro were cultured in SGal medium until reaching post-diauxic phase at 25°C, transferred to 37°C for 4 h and subjected to fluorescence microscopy (B,D). Cells were incubated for 30 mins with Mitotracker prior microscopy to stain the mitochondria (B). Quantification of Mitotracker fluorescent signal from at least 150 cells (C). Quantification of cells presenting 0, 1 or 2 Tim50 aggregates at 37°C is shown (E). (F) Wild-type, *tif51A-1* and *tif51A-1* carrying Tim50Pro were cultured as in (B). mRNA relative levels from mitoCPR genes were determined by RT-qPCR. (G) Wild-type, *tif51A-1*, wild-type Tim50Pro and *tif51A-1* Tim50Pro strains expressing Cyc1-nLuc (left) or Cox5a-nLuc (right) were cultured in YPD until early exponential phase and then transferred to 25°C or 37°C for 4 h. After addition of doxycycline to induce luciferase expression, the luminescence levels generated by the nanoluciferase after the addition of the furimazine substrate were measured along time and protein was quantified after 60 min. Data information: In (C,F,G) Results are presented as mean ± SD from three independent experiments. The statistical significance was measured by using a two-tailed Student t-test. *p < 0.05, **p < 0.001, ***p < 0.001.

We first tested the effect of substituting the endogenous Tim50 by Tim50Pro, whose translation is eIF5A-independent, on mitochondrial function and growth on respiratory media in eIF5A mutant cells. Expression of Tim50Pro did not rescue the growth of the *tif51A-1* mutant under glycerol at semi-permissive temperature (Fig 5A). This result points to the idea that Tim50 is not the only mechanism linking eIF5A to mitochondrial function. In addition, when we checked the accumulation of the membrane potential-dependent dye mitotracker red, we observed a slight increase in the membrane potential at permissive and restrictive temperatures in the *tif51A-1* Tim50Pro mutant respect to *tif51A-1* mutant cells expressing wild-type Tim50 (Figs 5B,C).

We then tested whether rescue of Tim50 translational stalling and protein levels would rescue the mitochondrial import phenotypes we observed in the eIF5A depletion. We found a reduction in Tim50 cytosolic protein aggregates upon removal of the stretch of prolines, indicating proper protein import (Figs 5D,E). Furthermore, we found that the mRNA levels of the most induced mitoCPR genes (*PDR3, PDR5* and *CIS1*) were significantly decreased in the *tif51A-1* Tim50Pro mutant (Fig 5F). As Tim50Pro rescued both the mitoCPR effect and protein import of Tim50, we next tested whether this may influence the translation of oxidative phosphorylation reporters that have reduced expression upon eIF5A depletion in cells expressing wild-type Tim50. We observed that eIF5A depletion does not significantly affect the synthesis of Cyc1 and Cox5A reporters if the translational stalling along *TIM50* is rescued by removing the prolines (Fig 5G). This suggests that the translational defects observed upon eIF5A depletion are secondary effects driven by a mitochondrial import stress.

## Discussion

In this study we have expanded our knowledge on the effects of eIF5A depletion on the mitochondrial function and identified one of the possible molecular mechanisms by which eIF5A is required to maintain mitochondrial activity. We have shown that eIF5A is necessary for the translation of the proline-rich region of Tim50 protein, which is part of the TIM23 complex and essential for the recognition and sorting of mitoproteins into the mitochondrial inner membrane and matrix. Tim50 and Tim23 proteins expose conserved soluble domains into the IMS that interact with each other, and this interaction is essential for protein translocation by the complex (53). Through the alleviation of ribosome stalling of *TIM50* mRNA on the mitochondrial surface and thus, modulation of Tim50 protein levels, eIF5A specifically impacts mitochondrial import. Upon eIF5A depletion, the mitochondrial import of many inner mitoproteins is compromised. We have seen that non-imported mitoproteins aggregate in the cytosol as a consequence of the import stress response provoked by the lack of eIF5A. Interestingly, it has been recently demonstrated that decreased mitoprotein uptake with CCCP causes the stall of translocating proteins in the outer membrane (54). A response for the clearance of stalled proteins in the mitochondrial surface has been shown to be mediated by the mitoCPR response, which induces, through the transcription factor Pdr3, the coordinated action of Cis1 and Msp1 to promote the degradation of arrested precursor proteins (18). Altogether, our results demonstrate that low levels of eIF5A causes mitochondrial stress by inducing *TIM50* ribosome stalling and reducing Tim50 expression (Fig 6).

**Fig. 6.**
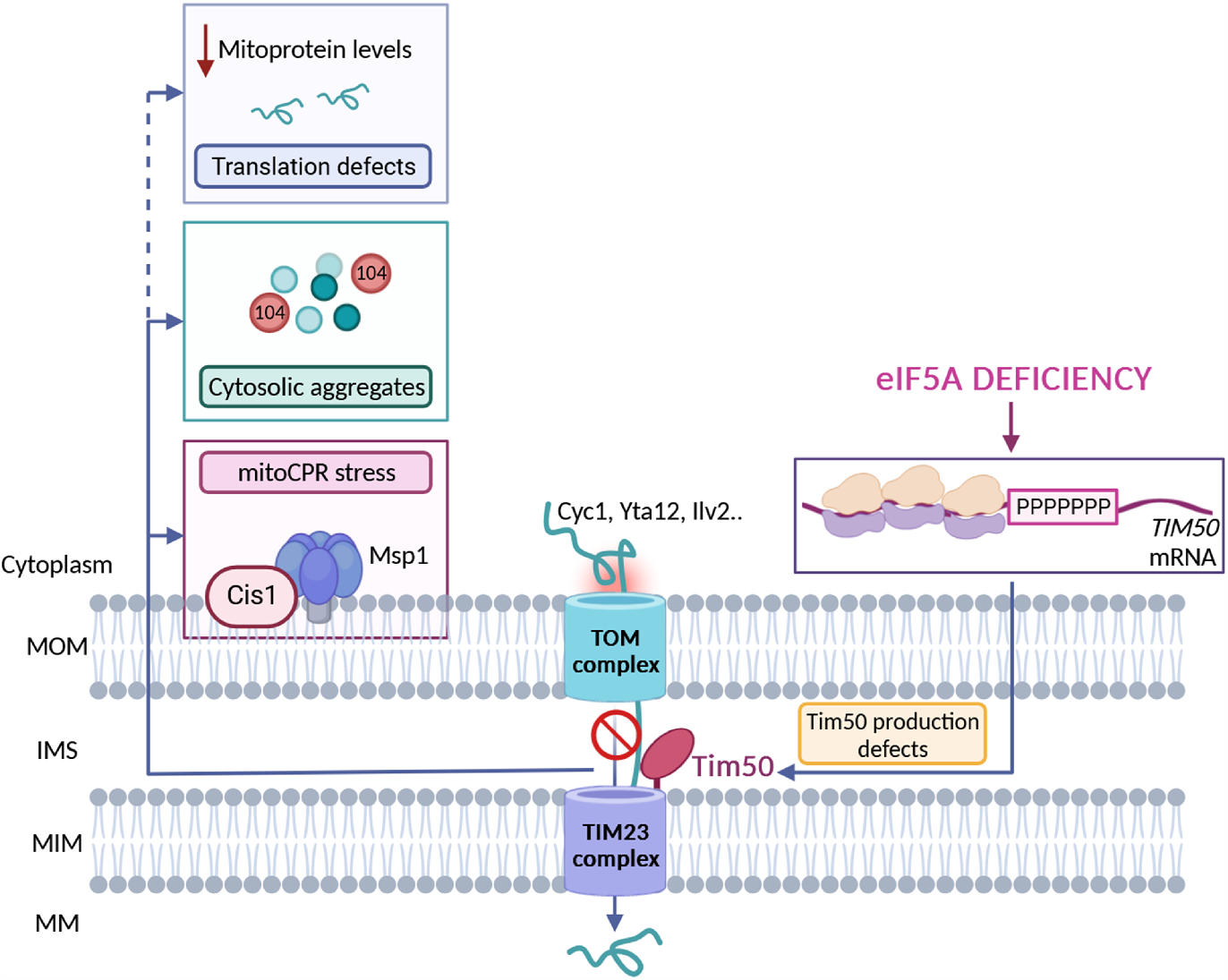
Model for the eIF5A-mediated regulation of the mitochondrial function. Upon eIF5A depletion, ribosomes translating TIM50 mRNA to synthesize the Tim50 proline-rich region become stalled and, thus, the mitochondrial import of Tim50-dependent proteins targeting the IMS, MIM and MM, is compromised. The non-imported mitochondrial precursors start aggregating in the cytosol bound to chaperone Hsp104 and the proteins translocating in the outer membrane become stalled. Then, the mitoCPR response is induced to clear the proteins accumulating in the mitochondrial surface through the action of Cis1 and Msp1. In addition, the lack of eIF5A reduces the translation of many RNAs encoding mitoproteins so the levels of most mitochondrial proteins become reduced. MOM: mitochondrial outer membrane; IMS: intermembrane space; MIM: mitochondrial inner membrane; MM: mitochondrial matrix. Figure was processed using BioRender software.

Our results also indicate that the accumulation of non-imported precursors in specific deposits in the cytosol is buffered by cytosolic chaperones, mainly Hsp104, to relieve the proteotoxic stress. These results are in line with the recent description of the cytosol as a place with capacity to store mitochondrial precursor proteins in dedicated storage granules that are controlled by the cytosolic chaperone system, with Hsp104 binding the N-terminal presequences of mitoproteins (41). Similarly, the observed Pdr5 induction upon eIF5A depletion seems indicative of a cellular detoxification effort to eliminate toxic substrates accumulating in the cytosol in a context of protein aggregates accumulation.

Upon prolonged mitochondrial dysfunction, the stress response is usually accompanied by cytosolic translation attenuation to reduce the synthesis of precursors and the protein load to translocases. General translation is known to be down-regulated upon treatment with multiple mitochondrial stressors such as defective mitochondrial biogenesis (16), clogger expression (17), oxidative stress (**?**) and mitochondrial depolarization with high CCCP doses (54). Herein, we have found that eIF5A depletion specifically down-regulates the expression of most mitoproteins. We observed that while general cytosolic translation remained unaffected, translation of proteins targeting mitochondria was found to be uniformly affected, suggesting a connection between the import status and the translation of mitoproteins that need to be translocated. In the last years it has been proposed that eIF5A is necessary for the translation of specific mitoproteins with that specificity residing in the amino acid sequences at their N-terminal/MTS sequences, especially those ones with weak interactions with the peptide exit tunnel, yet it was unclear if this was a direct effect (35, 36). The fact that Tim50Pro alleviated the translational repression of Cyc1 and Cox5A even upon eIF5A depletion suggests that some aspects of this translational regulation may result indirectly from eIF5A depletion. This could potentially be linked to the mitochondrial import stress response rather than being directly regulated by eIF5A. Nevertheless, more work will be needed to further decipher the specific molecular changes linking eIF5A, general translation of mitochondrial proteins and overall mitochondrial function.

The presented results herein highlight the idea that multiple mechanisms, besides Tim50 regulation, link eIF5A to the mitochondrial function. While we found the recovery of Tim50 function with the Tim50Pro to rescue the mitochondrial import stress response after 4 hours of eIF5A depletion, this was not sufficient to fully recover the MIM potential nor to rescue the growth on respiratory media for longer times at semi-permissive temperature. This could be explained by the fact that over sustained periods at semi-permissive temperatures, eIF5A becomes further depleted and causes other mitochondrial mRNAs that are less sensitive to eIF5A levels to become stalled and to drive similar mitochondrial stress responses. Therefore, it is of interest to explore other putative mitochondrial targets with eIF5A-dependent motifs in their sequences.

Human Tim50 sequence contains 27 prolines, but these are found scattered throughout the sequence rather than accumulated in a specific region as in *S. cerevisiae’s* Tim50. In future studies, it would be of interest to examine if Tim50 is a true eIF5A-target for its translation in higher eukaryotes. In humans, Tim50 down-regulation and derived import failure is associated with neurodegeneration and several genetic disorders (55, 56) whereas Tim50 overexpression has been observed in some cancer cell lines. The up-regulation of Tim50 activity may increase protein import and mitochondrial function, promoting cell growth and metastasis (57–59). Furthermore, clogging of translocases and cytosolic deposition of mitochondrial precursors have been implicated with some neurodegenerative disorders. Aggregation of mitoproteins in the cytosol increases misfolding of α-synuclein and amyloid precursor proteins, which are involved in Parkinson’s and Alzheimer’s diseases, as they can co-aggregate together, engage translocases and further increase the clogging hazard (40, 60). The specific eIF5A-mediated mechanism of Tim50 translational stalling alleviation and therefore, mitochondrial protein import regulation, might be important for understanding the molecular basis of pathologies where mitochondrial protein uptake fails. Importantly, data shown herein points to a novel and more likely mitochondrial stress generated by lack or defects in eIF5A protein that could be found in nature and raise as one of the underlying causes of impaired protein uptake in disease contexts. Thereby, eIF5A and its well-characterized hypusination precursor spermidine, could be considered as potential candidates to potentiate the activity of mitochondrial import machineries in compromised cells.

## Materials and methods

### Yeast strains, plasmids, and growth conditions

All *Saccharomyces cerevisiae* strains and plasmids used herein are listed in Supplementary Tables S2 and S3 respectively. *S. cerevisiae* cells were grown in either liquid YPD (2% glucose, 2% peptone, 1% yeast extract), YPGly (2% glycerol, 2% peptone, 1% yeast extract), synthetic complete medium (SC) [2% glucose, 0.7% yeast nitrogen base (YNB) and Drop-Out complete (Kaiser, Formedium)] or SGal (2% galactose, 0.7% YNB and Drop-Out complete).

For protein visualization under the microscope, a PCR-based genomic tagging technique was employed to tag the genomic full length *TIM50, TOM70, CYC1, YTA12, ILV2* and *PDR5* ORFs with GFP or the *HSP104* ORF with RFP. The plasmids pFA6a-GFP-HIS3MX (61) and pYM43-Redstar2-clonNAT were used as a template for PCR reaction using primers listed in Table S4. The plasmid pFA6a-3HA-HIS3MX (Longtine et al, 1998) was used to tag the genomic full length *TIM50* using primers listed in Table S4. The resulting cassettes were transformed in the corresponding strains following the lithium acetate-based method (62). All the integrations were confirmed by genomic DNA conventional PCR.

Plasmid pRS406-GPDp-Su9-mCherry-URA3 was integrated in the corresponding strains for expression of Su9 (subunit 9 of the F0 ATPase) fused to mCherry and subsequent visualization of mitochondrial network. To constitutively visualize *TIM50* single mRNAs, plasmids pRS405-CYC1p-MS2-4xGFP-LEU2 and plasmid pRS403-TIM50p-TIM50-flagiRFP-TIM50ter-MS2tag-HIS3MX (10) were integrated in wild-type and *tif51A-1* strains.

Plasmid pGAL-FLAG-TIM50-GFP-URA3 was constructed from the plasmid pYES2-pGAL-FLAG-htt25QP-GFP-URA3 (63). First, the yeast *TIM50* ORF was obtained by PCR using primers listed in Table S4. Then, plasmid pYES2-pGAL-FLAG-htt25QP-GFP-URA3 was linearized by restriction enzyme XagI (Thermo Fisher Scientific) to replace huntingtin (htt) gene by *TIM50* gene by homologous recombination. The resulting plasmid was transformed in the corresponding strains and transformants were selected in SC medium lacking uracil. To generate the strains harbouring the deletion of the seven consecutive prolines from the *TIM50* gene sequence (Tim50Pro), the C-terminal TIM50-GFP sequence was amplified from genomic DNA of the strain PAY1078 using primers listed in Table S4. The use of these primers resulted in the deletion of nucleotides 541-561 in *TIM50*, which encode for the 7 prolines stretch of the Tim50 protein. The resulting PCR product was transformed in wild-type and *tif51A-1* strains as previously described. All the integrations and deletions were confirmed by genomic DNA conventional PCR.

Nanoluciferase reporter constructs expressing different versions of the *TIM50* gene or other mitoproteins under the control of a tetracycline-inducible operon (tetO7) were generated by cloning the corresponding ORF in the plasmid ZP446, derived from pAG306 vector. ZP446 backbone was amplified by PCR using oligos listed in Table S4. Parental plasmid was digested by restriction enzyme DpnI (New England Biolabs) and linearized plasmid was isolated by gel purification (Zymo Research). The CDS fragments for cloning were amplified by conventional PCR from wild-type genomic DNA using oligos listed in Table S4. After PCR product purification (Zymo Research), the fragments of interest were inserted into linearized ZP446 using Gibson Assembly. The resulting nanoluciferase constructs were linearized by restriction enzyme NotI (New England Biolabs) and integrated into the genome of the wild-type and *TIM50* cells by homologous recombination. All the Tim50-nLuc plasmids used in this study are listed in Table S3.

Experimental assays were performed with cells exponentially grown for at least four generations until required OD600 at the corresponding temperature. Temperature-sensitive strains were grown at the permissive temperature of 25°C until required OD600 and transferred to the non-permissive temperature of 37°C or 41°C for 4 h for complete depletion of eIF5A.

### Proteomic analysis

For the proteomic analysis three independent cultures of wild-type (BY4741), *tif51A-1* and *tif51A-3* were grown exponentially in YPD at 25°C and then incubated at 41°C for 4h. Proteins were extracted as previously described (38) and analysed in the SCSIE (Servei Central de Suport a la Investigació Experimental; Universitat de València). Protein samples (about 20 μg of protein) were digested with 500 ng of trypsin (Promega) and peptides were analysed by an Ekspert nanoLC 42 nanoflow system (Eksigent Technologies, ABSCIEX) coupled to a nanoESI qQTOF MS (6600plus TripleTOF, ABSCIEX). The tripleTOF was operated in SWATH mode. 5 μl of each sample were loaded onto a trap column (3 μ C18-CL 120 Ă, 350 μm x 0.5 mm; Eksigent) and desalted with 0.1% TFA at 5 μl/min during 5 min. The peptides were loaded onto an analytical column (3 μ C18-CL 120 Ă, 0.075 × 150 mm; Eksigent) equilibrated in 5% acetonitrile 0.1% FA (formic acid). Peptide elution was carried out with a linear gradient of 7 to 40% B in 45 min (A: 0.1% FA; B: ACN, 0.1% FA) for at a flow rate of 300 nl/min. Peptides were analysed in a mass spectrometer nanoESI qQTOF (6600plus TripleTOF, ABSCIEX) using positive electrospray ionization (ESI) at an ion source temperature of 200°C. The tripleTOF was operated in swath mode, in which a 0.050-s TOF MS scan from 350–1250 m/z was performed, followed by 0.080-s product ion scans from 350–1250 m/z. 100 variable windows from 400 to 1250 m/z were acquired throughout the experiment. The total cycle time was 2.79 secs.

The processing settings used for the peptide selection were: detect at least 20 peptides per protein, 6 or more transitions per peptide, 95% peptide confidence threshold and 1% false discovery rate threshold; modified peptides were excluded. Only proteins that met these criteria and were detected in all strains were analysed in this study.

To analyse the proteomic data, protein areas were normalized by the total sum of the areas of all the quantified proteins. To obtain the GO terms overrepresented in the different groups of proteins, the ratio 41°C/25°C was calculated for each replicate and strain. Then each 41°C/25°C ratio of each temperature-sensitive mutant was divided by the ratio 41°C/25°C of wild-type strain, so that those with a value greater than 1 are up-regulated in the temperature-sensitive mutant with respect to wild-type, while those with a value less than 1 are down-regulated. Statistical significance was measured by Student’s t-test; only statistically significant proteins (p-value Student’s t-test < 0.05) were analysed.

### Nanoluciferase reporter assays

Nanoluciferase synthesis assays were performed as detailed in (64). Briefly, cells were grown at 25°C in YPD media to an exponential OD600 0.4 and then transferred to 37°C for 4 h. Doxycycline was added to a final concentration of 10 μg/mL to induce the expression of the nanoluciferase reporter of the pAG306 series vectors and pre-incubated for 5 min at room temperature. The nanoluciferase activity was measured using furimazine as the nanoluciferase highly specific substrate. A 90 μl volume of each culture and 10 μl of the furimazine (1 in 200 dilution; Promega) were incubated in a Cellstar non-transparent white 96-well microplate for one h and the bioluminescence intensity was monitored with a Tecan Infinite 200 PRO plate reader every 30 s. All the bioluminescence measurements were acquired at 460 nm, the peak emission wavelength of nanoluciferase. Obtained data were linearized using Schleif plots to estimate the minimum reaction time required for complete translation (50). The reaction time of the nLuc reporter alone was subtracted from the reaction time of the corresponding Tim50-nLuc fusion to calculate the time required for translating the different Tim50 sequence versions. At least five biological replicates of each Tim50-nLuc construct were analysed.

### RT-qPCR analysis

For the analysis of the mRNA levels, total RNAs were isolated from yeast cells following the phenol:chloroform protocol. Briefly, a volume of an exponential phase culture corresponding to 10 OD600 units was harvested and flash frozen. Cells were resuspended in 500 μL of cold LETS buffer (LiCl 0.1 M, EDTA pH 8.0 10 mM, Tris-HCl pH 7.4 10 mM, SDS 0.2%) and transferred into a screw-cap tube already containing 500 μL of sterile glass beads and 500 μL of phenol:chloroform (5:1). Then, cells were broken using the Precellys 24 tissue homogenizer (Bertin Technologies) and centrifugated. The supernatant was transferred into a new tube containing 500 μL of phenol:chloroform (5:1) and then to a tube containing 500 μL of chloroform:isoamyl alcohol (25:1). RNA from the top phase was precipitated and finally dissolved in water for later quantification and quality control with Nanodrop device (Thermo Fisher Scientific).

The reverse transcription and quantitative PCR reactions were performed as detailed in (65). Briefly, 2.5 μg of the total DNAse-I (Roche) treated RNA were retrotranscribed using an oligo d(T)18 with Maxima Reverse Transcriptase (Thermo Fisher Scientific). cDNA was labelled with SYBR Pre-mix Ex Taq (Tli RNase H Plus, Takara) and the Cq values were obtained from the CFX96 TouchTM Real-Time PCR Detection System (BioRad). Endogenous *ACT1* mRNA levels were used for normalization. At least three biological replicates of each sample were analysed, and the specific primers designed to amplify gene fragments of interest are listed in Table S4.

### Western blotting

For yeast protein content analysis by western blotting we followed the protocol described in (66). Briefly, a cell culture volume corresponding to 10 OD600 units was harvested by centrifugation. For protein extraction, cell pellets were washed and resuspended in 200 μL of NaOH 0.2M and incubated at room temperature for 5 min for subsequent centrifugation at 12000 rpm for 1 min. Samples were then resuspended in 100 μL of 2X-SDS protein loading buffer (24 mM Tris-HCl pH 6.8, 10% glycerol, 0.8% SDS, 5.76 mM β-mercaptoethanol, 0.04% bromophenol blue) and boiled at 95°C for 5 min. Next, lysates were centrifuged at 3000 rpm for 10 min at 4°C to remove cell debris and insoluble proteins, and supernatants were transferred into new tubes and stored at -20°C. Total protein content in the extract was quantified by an OD280 estimation in a Nanodrop device (Thermo Fisher Scientific) to load equal protein amounts per sample into the SDS-PAGE gel. The acrylamide percentage of the used SDS-PAGE depended on the molecular weight of the protein of interest. SDS-PAGE and Western blotting were performed using standard procedures (BioRad). Blotting membranes were blocked with 5% skimmed milk in TBS-T (150 mM NaCl, 20 mM Tris, 0.1% Tween20, pH 7.6) for 1 h at room temperature and incubated with primary antibodies overnight at 4°C against either HA (1:5000, Roche 12013819001), FLAG (1:1000, Sigma F1804), Por1 (1:1000, Abcam ab110326), Hsp60 (1:1000, QED-BIOSCIENCE 11101) or glyceraldehyde-6-phosphate dehydrogenase (rabbit polyclonal anti-G6PDH antibody, 1:10000, Sigma A9521). Bound antibodies were detected using the appropriate horseradish peroxidase-conjugated secondary antibodies (1:10000, Promega). Chemiluminiscent signals were detected with an ECL Prime Western blotting detection kit (GE Healthcare) and digitally analysed using ImageQuant LAS 4000 software (GE Healthcare). Intensity of bands was normalized against G6PDH bands. At least three biological replicates of each sample were analysed.

### Fluorescence microscopy and analysis

Yeast cells were grown to a post-diauxic phase in SGal medium, centrifugated, washed and subjected to standard fluorescence and phase contrast microscopy. For mitochondrial protein localization experiments, cells were imaged using an Eclipse Ti-E microscope (Nikon) with an oil-immersion x63 objective. Imaging was controlled using NIS-Elements software (Nikon). For mitochondrial aggregates co-localization experiments and mitochondrial membrane potential measurements, fluorescence images were acquired using an Axio Imager Z1 fluorescence motorized microscope equipped with a Plan Apochratic x63/1.4 oil-immersion objective and a 100W mercury lamp (Carl Zeiss, Germany). Images were recorded with an AxioCam MRm digital camera (Carl Zeiss, Germany). The following excitation and emission wavelengths were used: DAPI (excitation 359 nm; emission 457 nm), GFP (excitation 475 nm; emission 509 nm), MitoTracker Red (excitation 578 nm; emission 600 nm), RFP (excitation 555 nm; emission 583 nm), mCherry (excitation 587 nm; emission 610 nm) and Nile Red (excitation 460 nm; emission 582 nm). The same exposure times were used to acquire all images. All the imaging analysis was performed on Image J software.

To analyse mitochondrial membrane potential, cells were incubated with 0.5 μM MitoTracker Red CMXRos (Thermo Fisher Scientific) for 30 min, washed and subjected to microscope. To analyse Pdr5 activity, cells were incubated with 3.5 μM Nile Red (Thermo Fisher Scientific) for 15 min, washed and subjected to microscope. To study nuclei localization, cells were incubated with 1 μg/mL 4,6-Diamidino-2-phenylindole dihydrochloride (DAPI, Thermo Fisher Scientific) for 20 min in the dark, washed and subjected to microscope.

For single molecule mRNA visualization with mitochondria, cells were imaged by an Eclipse Ti2-E Spinning Disk Confocal with Yokogawa CSU-X1 (Yokogawa) with 50 μm pinholes, located at the Nikon Imaging Center UC San Diego. Imaging was performed using SR HP APO TIRF 100 × 1.49 NA oil objective with the correction collar set manually for each experiment (pixel size 0.074 mm). Z-stacks (300 nm steps) were acquired by a Prime 95B sCMOS camera (Photometrics). Imaging was controlled using NIS-Elements software (Nikon).

### Polyribosome profile analysis

For polysome fractioning, cells were grown at 25°C to post-diauxic phase in SGal medium and transferred to 37°C for 4 h for the experiments requiring temperature-sensitive strains. Cell extractions and polysome gradients were performed as described by (67). Briefly, a culture volume corresponding to an OD_600_ of 100 was chilled for 5 min on ice in the presence of 0.1 mg/mL CHX. Cells were centrifuged at 4400 rpm for 3 min at 4°C and washed twice with 2 mL of lysis buffer (20 mM Tris-HCl pH 8, 140 mM KCl, 5 mM MgCl_2_, 0.5 mM DTT, 1% Triton X-100, 0.1 mg/mL CHX, and 0.5 mg/mL heparin). Cells were resuspended in 700 μL of lysis buffer and added to a tube already containing 500 μL of glass beads. Cells were mechanically disrupted by vortexing 8 times for 30 s with 30 s of incubation on ice in between. Then, lysates were cleared by centrifugation at 5000 rpm for 5 min at 4°C, and the supernatant was recovered. After centrifugation at 8000 rpm for 5 min at 4°C, the RNA was recovered, and its concentration was estimated using a Nanodrop device (Thermo Fisher Scientific). Glycerol was added to all the samples to a final concentration of 5%, and extracts were flash-frozen and stored at -80°C. Samples of 10 A_260_nm units were loaded onto 5-50% sucrose gradients and separated by ultracentrifugation for 2 h 40 min at 3500 rpm in a Beckman SW41Ti rotor at 4°C. Then, gradients were fractionated by isotonic pumping of 60% sucrose from the bottom, and twenty-two 0.5 mL samples were recovered. The polysome profiles were monitored by UV detection at 260 nm using a density gradient fractionation system (Teledyne Isco, Lincoln, NE). RNAs were extracted using SpeedTools Total RNA Extraction kit (Biotools B&M Labs) with the rDNAse treatment after the RNA elution step. Specific mRNAs were analyzed by RT-qPCR using specific primers (listed in Table S4) and represented as a percentage of total. Three biological replicates were performed for each polyribosome profile, and a representative profile is shown.

### Proteomic data accession

The mass spectrometry proteomics data have been deposited to the ProteomeXchange Consortium (68) via the PRIDE (69) partner repository with the dataset identifier PXD043905.

## Acknowledgements

We thank Oreto Antúnez and M. Luz Valero from the Proteomics Service SCSIE (Universitat de València). Grant PID2020-120066RB-I00 funded by MCIN/AEI/10.13039/501100011033 to PA. This work was supported, in part, by the National Institutes of Health R35GM128798 to BMZ. This research was also funded by Generalitat Valenciana (AICO/2020/086 and CIAICO/2022/237) to PA. MB-A. is a recipient of a predoctoral fellowship (FPU2017/03542) funded by MCIN/AEI/10.13039/501100011033 and by ESF Investing in your future. Authors acknowledge support by all members of GFL lab.

## Author contributions

PA, BMZ and MB-A conceived and supervised the study. PA, BMZ and MB-A designed the experiments. MB-A, VB and CR performed the experiments. MB-A PA and BMZ wrote and revised the manuscript.

## References

1. AJ Roger, SA Munoz-Gomez, and R Kamikawa. The origin and diversification of mitochondria. Curr Biol, 27:1177–1192, 2017.

2. Robert S Balaban, Shumpei Nemoto, and Toren Finkel. Mitochondria, oxidants, and aging. Cell, 120:483–495, 2005.

3. Kerstin Boengler, Miriam Kosiol, Manuel Mayr, Rainer Schulz, and Susanne Rohrbach. Mitochondria and ageing: role in heart, skeletal muscle and adipose tissue. Journal of Cachexia, Sarcopenia and Muscle, 8:349–369, 2017.

4. O Schmidt, N Pfanner, and C Meisinger. Mitochondrial protein import: From proteomics to functional mechanisms. Nat Rev Mol Cell Biol, 11:655–667, 2010.

5. Peter Borst and Leo A Grivell. Mitochondrial nucleic acids. Biochimie, 15:705–723, 1973.

6. Yury S Bykov, Doron Rapaport, Johannes M Herrmann, and Maya Schuldiner. Cytosolic events in the biogenesis of mitochondrial proteins. Trends in Biochemical Sciences, 45: 650–667, 2020.

7. Katja G Hansen and Johannes M Herrmann. Transport of proteins into mitochondria. Protein Journal, 38(4):330–342, 2019.

8. Rodney E Kellems and Ronald A Butow. Cytoplasmic-type 80 s ribosomes associated with yeast mitochondria. i. evidence for ribosome binding sites on yeast mitochondria. Journal of Biological Chemistry, 247(24):8043–8050, 1972.

9. CC Williams, CH Jan, and JS Weissman. Targeting and plasticity of mitochondrial proteins revealed by proximity-specific ribosome profiling. Science (80-), 346:748–751, 2014.

10. T Tsuboi, MP Viana, F Xu, J Yu, R Chanchani, XG Arceo, E Tutucci, J Choi, YS Chen, RH Singer, et al. Mitochondrial volume fraction and translation duration impact mitochondrial mrna localization and protein synthesis. Elife, 9:e57814, 2020.

11. FN Vögtle, S Wortelkamp, RP Zahedi, D Becker, C Leidhold, K Gevaert, J Kellermann, W Voos, A Sickmann, N Pfanner, et al. Global analysis of the mitochondrial n-proteome identifies a processing peptidase critical for protein stability. Cell, 139:428–439, 2009.

12. N Wiedemann and N Pfanner. Mitochondrial machineries for protein import and assembly. Annu Rev Biochem, 86:685–714, 2017.

13. Anna Maria Lenkiewicz, Michał Krakowczyk, and Piotr Bragoszewski. Cytosolic quality control of mitochondrial protein precursors—the early stages of the organelle biogenesis. International Journal of Molecular Sciences, 23(7), 2022.

14. Mai Oxholm Haastrup, Kailash Singh Vikramdeo, Shikha Singh, Anand Prakash Singh, and Santanu Dasgupta. The journey of mitochondrial protein import and the roadmap to follow. International Journal of Molecular Sciences, 24(5):2479, 2023.

15. X Wang and XJ Chen. A cytosolic network suppressing mitochondria-mediated proteostatic stress and cell death. Nature, 2015.

16. L Wrobel, U Topf, P Bragoszewski, S Wiese, ME Sztolsztener, S Oeljeklaus, A Varabyova, M Lirski, P Chroscicki, S Mroczek, et al. Mistargeted mitochondrial proteins activate a proteostatic response in the cytosol. Nature, 524:485–488, 2015.

17. Felix Boos, Lars Kramer, Christine Groh, Florian Jung, Per Haberkant, Ferris Stein, Florian Wollweber, Andreas Gackstatter, Eva Zöller, Martin van der Laan, et al. Mitochondrial protein-induced stress triggers a global adaptive transcriptional programme. Nature Cell Biology, 21:442–451, 2019.

18. H Weidberg and A Amon. Mitocpr—a surveillance pathway that protects mitochondria in response to protein import stress. Science (80-), 360:eaan4146, 2018.

19. Christoph U Mårtensson, Christian Priesnitz, Jiyao Song, Lars Ellenrieder, Kim N Doan, Franziska Boos, Anna Floerchinger, Nikolaus Zufall, Silke Oeljeklaus, Bettina Warscheid, et al. Mitochondrial protein translocation-associated degradation. Nature, 569(7757):679–683, 2019.

20. J Schnier, HG Schwelberger, Z Smit-McBride, HA Kang, and JW Hershey. Translation initiation factor 5a and its hypusine modification are essential for cell viability in the yeast saccharomyces cerevisiae. Mol Cell Biol, 11:3105–3114, 1991.

21. Emilio Gutierrez, Byung-Sik Shin, Christopher J Woolstenhulme, Joo-Ran Kim, Puneet Saini, Allen R Buskirk, and Thomas E Dever. eif5a promotes translation of polyproline motifs. Molecular Cell, 51(1):35–45, 2013.

22. Vicent Pelechano and Paula Alepuz. Eif5a facilitates translation termination globally and promotes the elongation of many non polyproline-specific tripeptide sequences. Nucleic Acids Research, 45(13):7326–7338, 2017.

23. AP Schuller, CCC Wu, TE Dever, AR Buskirk, and R Green. eif5a functions globally in translation elongation and termination. Mol Cell, 66:194–205, 2017.

24. Zhane A Jenkins, Petra G Hååg, and Hans E Johansson. Human eif5a2 on chromosome 3q25-q27 is a phylogenetically conserved vertebrate variant of eukaryotic translation initiation factor 5a with tissue-specific expression. Genomics, 71(1):101–109, 2001.

25. Ming-Hsu Park, Ronak K Kar, Siddharth Banka, Andreas Ziegler, and Wendy K Chung. Post-translational formation of hypusine in eif5a: implications in human neurodevelopment. Amino Acids, pages 1–15, 2021.

26. M Barba-Aliaga, C Villarroel-Vicente, A Stanciu, A Corman, MT Martínez-Pastor, and P Alepuz. Yeast translation elongation factor eif5a expression is regulated by nutrient availability through different signalling pathways. International Journal of Molecular Sciences, 22:219, 2021.

27. M Barba-Aliaga and P Alepuz. The activator/repressor hap1 binds to the yeast eif5aencoding gene tif51a to adapt its expression to the mitochondrial functional status. FEBS Letters, 596:1809–1826, 2022.

28. L Zhang and A Hach. Molecular mechanism of heme signaling in yeast: The transcriptional activator hap1 serves as the key mediator. Cell Mol Life Sci, 56:415–426, 1999.

29. M Barba-Aliaga and P Alepuz. Role of eif5a in mitochondrial function. International Journal of Molecular Sciences, 23:1284, 2022.

30. Nicolas Melis, Isabella Rubera, Marc Cougnon, Stephanie Giraud, Baharia Mograbi, Amine Belaid, Didier F Pisani, Stephan M Huber, Sandra Lacas-Gervais, Konstantina Fragaki, et al. Targeting eif5a hypusination prevents anoxic cell death through mitochondrial silencing and improves kidney transplant outcome. Journal of the American Society of Nephrology, 28(3): 811–822, 2017.

31. Stephanie Giraud, Thomas Kerforne, Julie Zely, Virginie Ameteau, Pascal Couturier, Michel Tauc, and Thierry Hauet. The inhibition of eif5a hypusination by gc7, a preconditioning protocol to prevent brain death-induced renal injuries in a preclinical porcine kidney transplantation model. American Journal of Transplantation, 20(12):3326–3340, 2020.

32. Jing Liu, Xianquan Zhan, Ming Li, Guan Li, Peng Zhang, Zhongde Xiao, Meichen Shao, Fuling Peng, Rui Hu, and Zhuchu Chen. Mitochondrial proteomics of nasopharyngeal carcinoma metastasis. BMC Medical Genomics, 5(1):1–17, 2012.

33. Takahito Miyake, Sunila Pradeep, Sherry Y Wu, Rajesha Rupaimoole, Behrouz Zand, Yunfei Wen, Kshipra M Gharpure, Archana S Nagaraja, Wei Hu, Mong S Cho, et al. Xpo1/crm1 inhibition causes antitumor effects by mitochondrial accumulation of eif5a. Clinical Cancer Research, 21(14):3286–3297, 2015.

34. KD Pereira, L Tamborlin, L Meneguello, ARG de Proenca, IC de PA Almeida, RF Lourenco, and AD Luchessi. Alternative start codon connects eif5a to mitochondria. J Cell Physiol, 231:2682–2689, 2016.

35. DJ Puleston, MD Buck, RI Klein Geltink, RL Kyle, G Caputa, D O’Sullivan, AM Cameron, A Castoldi, Y Musa, AM Kabat, et al. Polyamines and eif5a hypusination modulate mitochondrial respiration and macrophage activation. Cell Metab, 30:352–363, 2019.

36. Y Zhang, D Su, J Zhu, M Wang, Y Zhang, Q Fu, S Zhang, and H Lin. Oxygen level regulates n-terminal translation elongation of selected proteins through deoxyhypusine hydroxylation. Cell Rep, 39:110855, 2022.

37. Zhaohui Li, Franco J Vizeacoumar, Sara Bahr, Jie Li, Jonas Warringer, Frederick S Vizeacoumar, Renqiang Min, Benjamin Vandersluis, Jeremy Bellay, Michael Devit, et al. Systematic exploration of essential yeast gene function with temperature-sensitive mutants. Nature Biotechnology, 29(4):361–367, 2011.

38. Tengfei Li, Borja Belda-Palazón, Armando Ferrando, and Paula Alepuz. Fertility and polarized cell growth depends on eif5a for translation of polyproline-rich formins in saccharomyces cerevisiae. Genetics, 197(4):1191–1200, 2014.

39. Wanchen Hou, Veronica Harjono, Adam T Harvey, Arvind R Subramaniam, and Brian M Zid. Quantification of elongation stalls and impact on gene expression in yeast. RNA, 2023.

40. Urszula Nowicka, Piotr Chroscicki, Katrien Stroobants, Monika Sladowska, Mateusz Turek, Barbara Uszczynska-Ratajczak, Ritwick Kundra, Tomasz Goral, Michele Perni, Christopher M Dobson, et al. Cytosolic aggregation of mitochondrial proteins disrupts cellular homeostasis by stimulating the aggregation of other proteins. eLife, 10:e65484, 2021.

41. Lisa Krämer, Nicole Dalheimer, Markus Räschle, Zuzana Storchová, Jan Pielage, Franz Boos, and Johannes M Herrmann. Mitostores: chaperone-controlled protein granules store mitochondrial precursors in the cytosol. EMBO Journal, 42(9):e112309, 2023.

42. VPS Shakya, WA Barbeau, T Xiao, CS Knutson, MH Schuler, and AL Hughes. A nuclear-based quality control pathway for non-imported mitochondrial proteins. Elife, 10:e61230, 2021.

43. Y Sanchez and SL Lindquist. Hsp104 required for induced thermotolerance. Science (80-), 248:1112–1115, 1990.

44. Stephanie N Gates, Adam L Yokom, JiaBei Lin, Meredith E Jackrel, Alex N Rizo, Nathan M Kendsersky, Christopher E Buell, Elizabeth A Sweeny, Kristen L Mack, Eric Chuang, et al. Ratchet-like polypeptide translocation mechanism of the aaa+ disaggregase hsp104. Science, 357(6348):273–279, 2017.

45. Shannon M Hill, Sjoerd Hanzén, and Thomas Nyström. Restricted access: spatial sequestration of damaged proteins during stress and aging. EMBO Reports, 18(3):377–391, 2017.

46. Dejana Mokranjac and Walter Neupert. Thirty years of protein translocation into mitochondria: Unexpectedly complex and still puzzling. Biochimica et Biophysica Acta (BBA)-Molecular Cell Research, 1793(1):1400–1407, 2009.

47. Mousumi Chaudhuri, Abhay Tripathi, and Frank S Gonzalez. Diverse functions of tim50, a component of the mitochondrial inner membrane protein translocase. International Journal of Molecular Sciences, 23:7779, 2021.

48. Romain Christiano, Nandhini Nagaraj, Florian Fröhlich, and Tobias C Walther. Global proteome turnover analyses of the yeasts s.cerevisiae and s.pombe. Cell Reports, 9:1959–1965, 2014.

49. Anna E Masser, Ganapathi Kandasamy, Jayasankar M Kaimal, and Claes Andréasson. Luciferase nanoluc as a reporter for gene expression and protein levels in saccharomyces cerevisiae. Yeast, 33(5):191–200, 2016.

50. R Schleif, W Hess, S Finkelstein, and D Ellis. Induction kinetics of the l arabinose operon of escherichia coli. J Bacteriol, 115:9–14, 1973.

51. M Zhu, X Dai, and YP Wang. Real time determination of bacterial in vivo ribosome translation elongation speed based on laczα complementation system. Nucleic Acids Res, 44: e155–e155, 2016.

52. Jun Li and Bingdong Sha. The structure of tim50(164-361) suggests the mechanism by which tim50 receives mitochondrial presequences. Acta Crystallographica Section F: Structural Biology Communications, 71(9):1146–1151, 2015.

53. Lilit Gevorkyan-Airapetov, Keren Zohary, Dušan Popov-Č eleketić, Koyeli Mapa, Kai Hell, Walter Neupert, Abdussalam Azem, and Dejana Mokranjac. Interaction of tim23 with tim50 is essential for protein translocation by the mitochondrial tim23 complex. Journal of Biological Chemistry, 284(7):4865–4872, 2009.

54. JA Schäfer, S Bozkurt, JB Michaelis, K Klann, and C Münch. Global mitochondrial protein import proteomics reveal distinct regulation by translation and translocation machinery. Mol Cell, 82:435–446, 2022.

55. S Sugiyama, S Moritoh, Y Furukawa, T Mizuno, YM Lim, L Tsuda, and Y Nishida. Involvement of the mitochondrial protein translocator component tim50 in growth, cell proliferation and the modulation of respiration in drosophila. Genetics, 176:927–936, 2007.

56. MA Shahrour, O Staretz-Chacham, D Dayan, J Stephen, A Weech, N Damseh, H Pri Chen, S Edvardson, S Mazaheri, A Saada, et al. Mitochondrial epileptic encephalopathy, 3-methylglutaconic aciduria and variable complex v deficiency associated with timm50 mutations. Clin Genet, 91:690–696, 2017.

57. H Sankala, C Vaughan, J Wang, S Deb, and PR Graves. Upregulation of the mitochondrial transport protein, tim50, by mutant p53 contributes to cell growth and chemoresistance. Arch Biochem Biophys, 512:52–60, 2011.

58. Shou-Ping Gao, Hong-Feng Sun, Hui-Ling Jiang, Li-Dong Li, Xue Hu, Xing-E Xu, and Wen Jin. Loss of tim50 suppresses proliferation and induces apoptosis in breast cancer. Tumor Biology, 37:1279–1287, 2016.

59. X Zhang, S Han, H Zhou, L Cai, J Li, N Liu, Y Liu, L Wang, C Fan, A Li, et al. Timm50 promotes tumor progression via erk signaling and predicts poor prognosis of non-small cell lung cancer patients. Mol Carcinog, 39:110855, 2019.

60. Giovanna Cenini, Chris Rub, Marija Bruderek, and Wolfgang Voos. Amyloid β-peptides interfere with mitochondrial preprotein import competence by a coaggregation process. Molecular Biology of the Cell, 27:3257–3272, 2016.

61. Mark S Longtine, Arlene McKenzie, Douglas J Demarini, Neil G Shah, Achim Wach, Sophie Brachat, Peter Philippsen, and John R Pringle. Additional modules for versatile and economical pcr-based gene deletion and modification in saccharomyces cerevisiae. Yeast, 14 (10):953–961, 1998.

62. R. Daniel Gietz, Anne Stirling Jean, Richard A Woods, and Robert H Schiestl. Improved method for high efficiency transformation of intact yeast cells. Nucleic Acids Research, 20 (6):1425, 1992.

63. Lars L Berglund, Xin Hao, Bin Liu, Julie Grantham, and Thomas Nyström. Differential effects of soluble and aggregating polyq proteins on cytotoxicity and type-1 myosin-dependent endocytosis in yeast. Scientific Reports, 7:11328, 2017.

64. Andrew R Guzikowski, Adam T Harvey, Jun Zhang, Shisheng Zhu, Kelsey Begovich, Marla H Cohn, Jennifer E Wilhelm, and Brian M Zid. Differential translation elongation directs protein synthesis in response to acute glucose deprivation in yeast. RNA Biology, 19(5):636–649, 2022.

65. Elena Garre, Leticia Romero-Santacreu, Manuela Barneo-Muñoz, Alberto Miguel, José E Pérez-Ortín, and Pilar Alepuz. Nonsense-mediated mrna decay controls the changes in yeast ribosomal protein pre-mrnas levels upon osmotic stress. PLoS One, 8:e61240, 2013.

66. A Zuzuarregui, T Li, C Friedmann, G Ammerer, and P Alepuz. Msb2 is a ste11 membrane concentrator required for full activation of the hog pathway. Biochim Biophys Acta - Gene Regul Mech, 1849:722–730, 2015.

67. Elena Garre, Leticia Romero-Santacreu, Nikki De Clercq, Noelia Blasco-Angulo, Per Sunnerhagen, and Paula Alepuz. Yeast mrna cap-binding protein cbc1/sto1 is necessary for the rapid reprogramming of translation after hyperosmotic shock. Molecular Biology of the Cell, 23(1):137–150, 2012.

68. Eric W Deutsch, Nuno Bandeira, Yasset Perez-Riverol, Vishal Sharma, Jeremy J Carver, Luis Mendoza, Dipali J Kundu, Sheng Wang, Chaitanya Bandla, S Kamatchinathan, et al. The proteomexchange consortium at 10 years: 2023 update. Nucleic Acids Research, 51:D1539–D1548, 2023.

69. Y Perez-Riverol, J Bai, C Bandla, D Garcia-Seisdedos, S Hewapathirana, S Kamatchinathan, DJ Kundu, A Prakash, A Frericks-Zipper, M Eisenacher, et al. The pride database resources in 2022: A hub for mass spectrometry-based proteomics evidences. Nucleic Acids Res, 50:D543–D552, 2022.

