## Supplementary Figures for "eIF5A controls mitoprotein import by relieving ribosome stalling at the *TIM50* translocase mRNA"

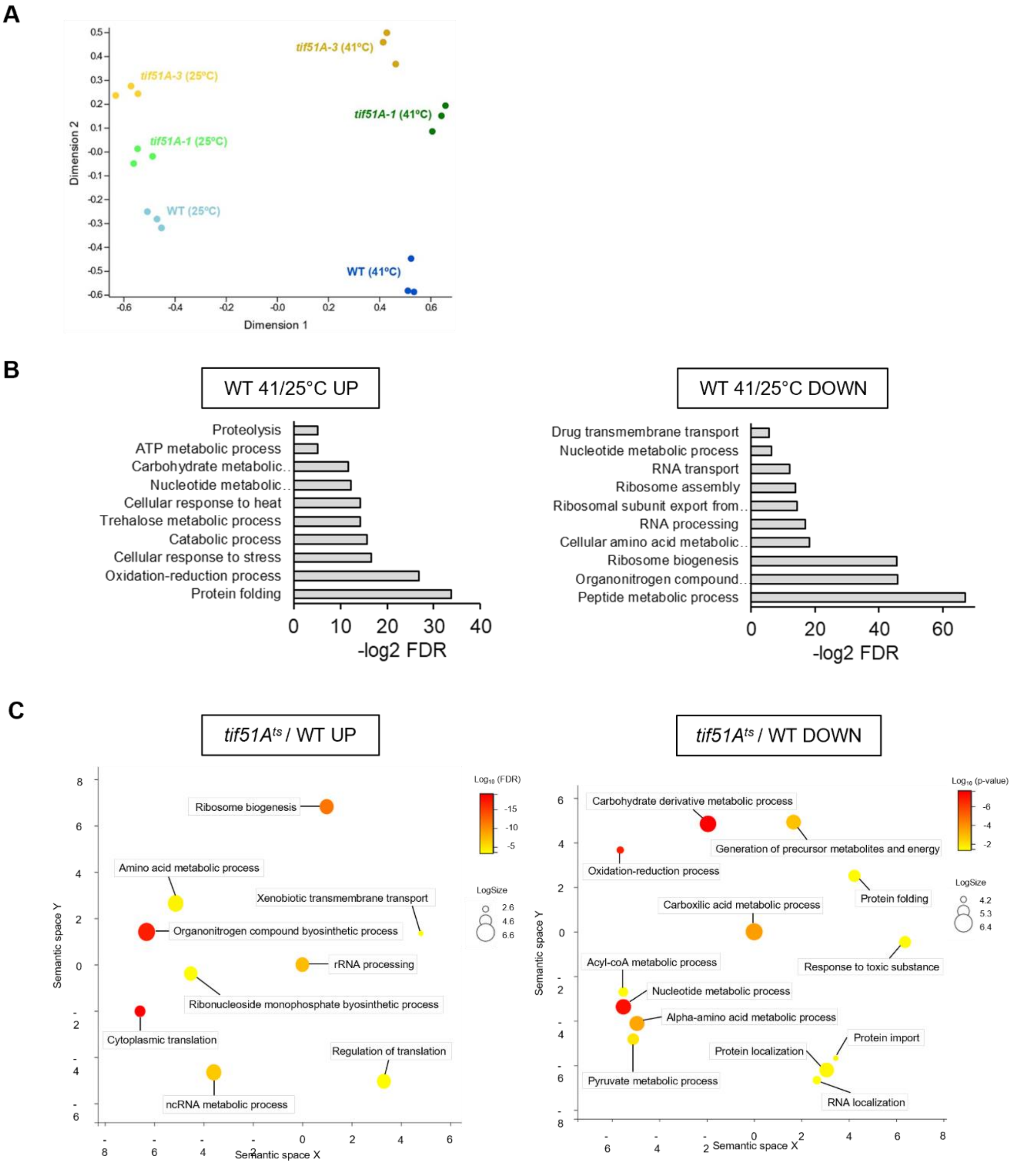


**Figure EV1. Proteomic analysis upon eIF5A depletion.**

(A) MDS-plot showing all replicates for each strain and condition studied in the proteomic analysis.

(B) Biological Process overrepresented in proteins up- or down- regulated significantly in wild-type cells at 41°C. GO Term Analysis was done using the STRING tool, and a total of 272 proteins up- (left) and 154 proteins down-regulated (right) significantly in WT 41°C compared to WT 25°C were analysed.

(C) Biological process Gene Ontology (GO) Terms overrepresented in relative up- (left) or down-regulated (right) proteins in *tif51A*$^{ts}$ vs WT. GO Term analysis was done using the STRING tool, in which a total of 292 proteins down- and 135 proteins up- regulated significantly in at least one *tif51A*$^{ts}$ with respect to WT were analysed. The web based tool REVIGO was used to summarize the GO terms. Bubble colour indicates the p-value of the GO term in the input data set; bubble size indicates the frequency of the GO term in the underlying GO database.

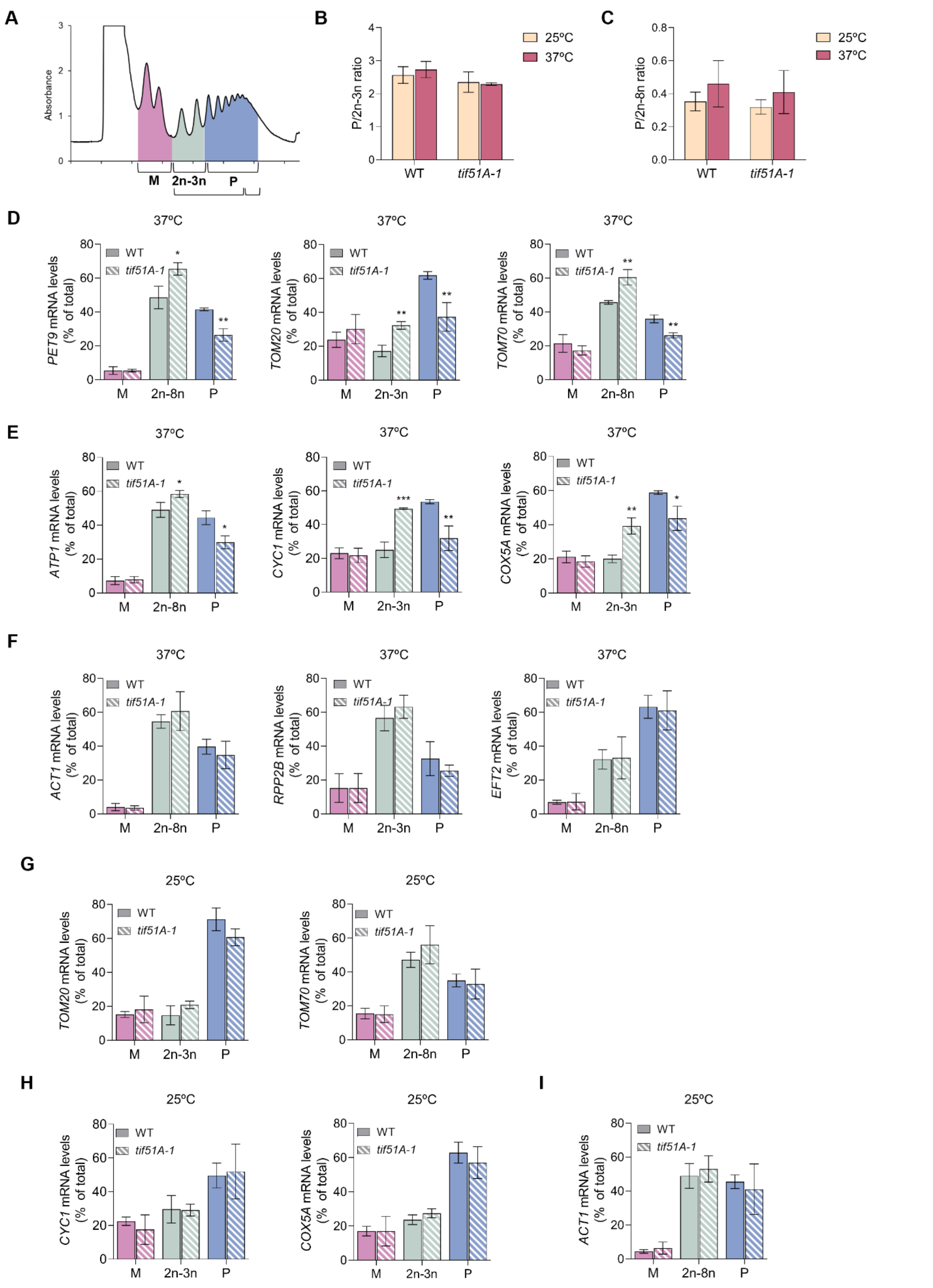


**Figure EV2. Translation of mitochondrial proteins is not affected in the *tif51A-1* strain at permissive temperature.**

(A) Scheme with the employed area divisions for calculations.

(B) The average polysomes/2n-3n ratio (P/2n-3n) is represented for each strain at both temperatures.

(C) The average polysomes/2n-8n ratio (P/2n-3n) is represented for each strain at both temperatures.

(D-F) The RNA from individual fractions of the polysomes profiles was extracted and the mRNA levels of *PET9*, *TOM20, TOM70* (D), *ATP1, CYC1, COX5A* (E), *ACT1, RPP2B* and *EFT2* (F) were analyzed by RT-qPCR in the corresponding sections at restrictive temperature. (G-I) The RNA from individual fractions of the polyribosome profiles was extracted and the mRNA levels of *TOM20*, *TOM70* (G), *CYC1*, COX5A (H) and *ACT1* (I) were analyzed by RT-qPCR in the corresponding sections at permissive temperature.

Data information: In (B-I) Results are presented as mean ± SD from three independent experiments. The statistical significance was measured by using a two-tailed paired Student t-test relative to wild-type strain. *$p < 0.05$, **$p < 0.001$, ***$p < 0.001$.

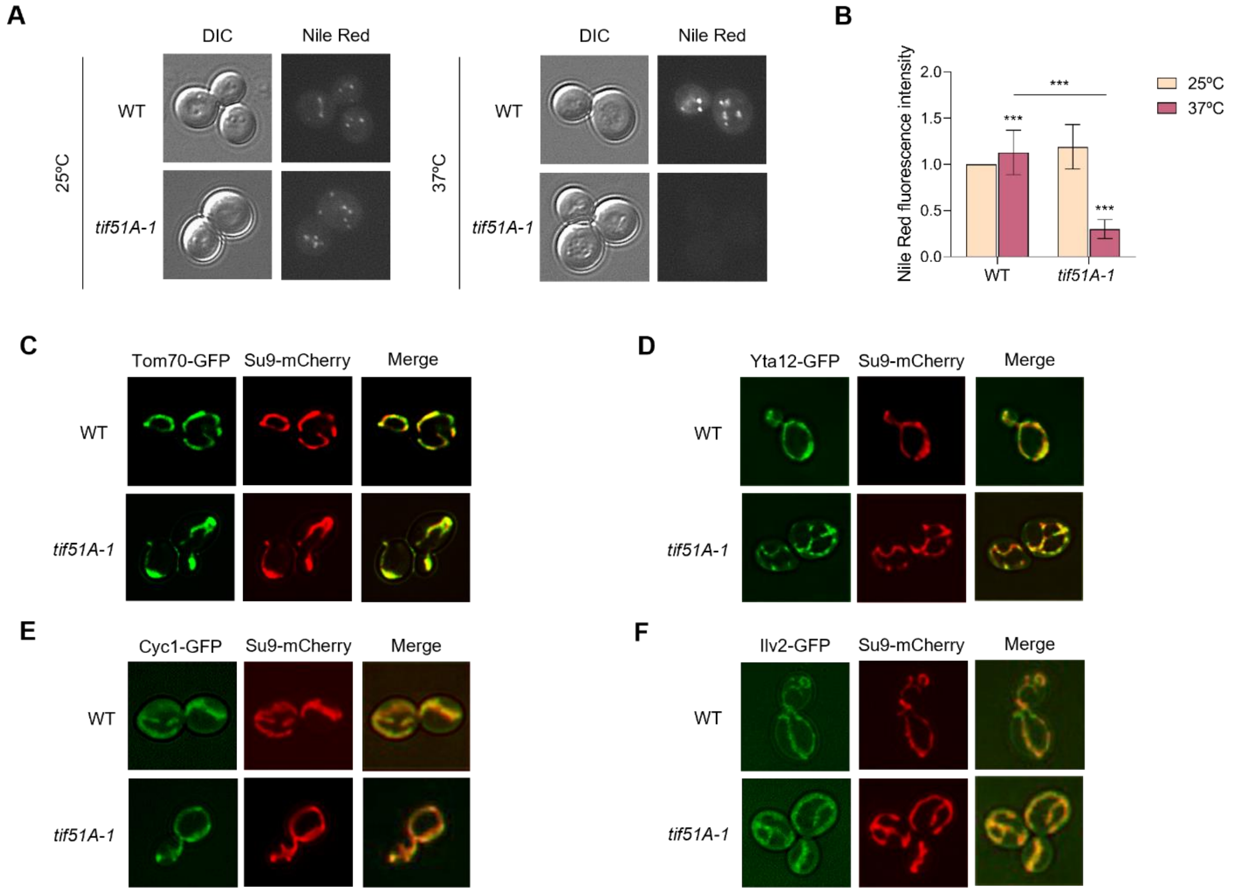


**Figure EV3. Nile Red is excluded from the *tif51A-1* mutant because of high Pdr5 pumping activity.**

(A) Wild-type strain and *tif51A-1* were cultured in SGal medium at 25°C until reaching post-diauxic phase and then transferred to 25°C or 37°C for 4 h. Then, cells were incubated with Nile Red substrate for 15 min prior to microscopy. A representative image is shown.

(B) Quantification of Nile Red fluorescent signal from at least 150 cells.

(C-F) Wild-type strain and *tif51A-1* expressing Tom70-GFP (C), Yta12-GFP (D), Cyc1-GFP (E) or Ilv2-GFP (F) and Su9-mCherry were cultured in SGal medium at 25°C until reaching post-diauxic phase and subjected to fluorescence microscopy.

Data information: In (B) Results are presented as mean ± SD from three independent experiments. The statistical significance was measured by using a two-tailed paired Student t-test relative to 25°C. ***$p < 0.001$.

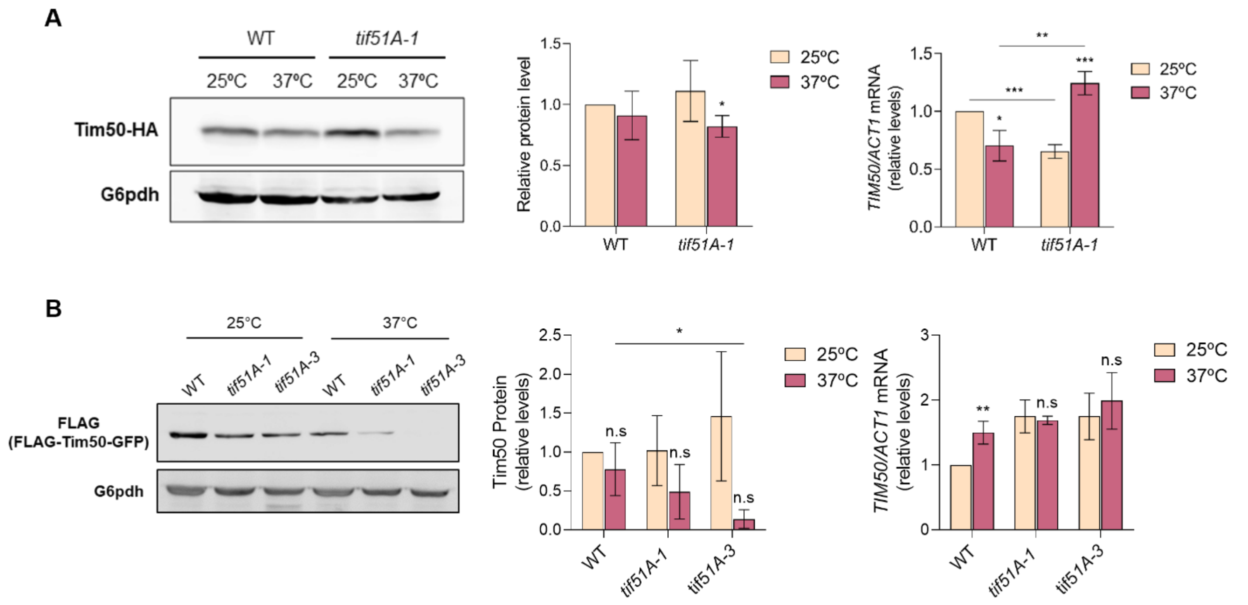


**Figure EV4. Tim50 protein levels are reduced upon eIF5A depletion.**

(A) Wild-type and *tif51A-1* strains containing genomic tagged Tim50-HA were cultured in SGal until post-diauxic phase at 25°C and transferred to 25°C or 37°C for 4 h. Tim50 protein levels were determined by western blotting (left) and quantified (middle). G6PDH levels were used as loading control. A representative image is shown. *TIM50* mRNA relative levels were determined by RT-qPCR (right).

(B) Wild-type, *tif51A-1* and *tif51A-3* strains harbouring a FLAG-TIM50-GFP plasmid were cultured in SRaf-URA at 25ºC until early exponential phase, transferred to 25°C and 37°C for 2 h and then transferred to SGal-URA at 25°C and 37°C for 3 additional hours. Tim50 protein levels were determined by western blotting using a FLAG antibody (left) and quantified (middle). G6PDH levels were used as loading control. A representative image is shown. *TIM50* mRNA relative levels were determined by RT-qPCR (right).

Data information: In (A,B) Results are presented as mean ± SD from three independent experiments. The statistical significance was measured by using a two-tailed paired Student t-test relative to 25ºC. *p < 0.05, **p < 0.01, ***p < 0.001.

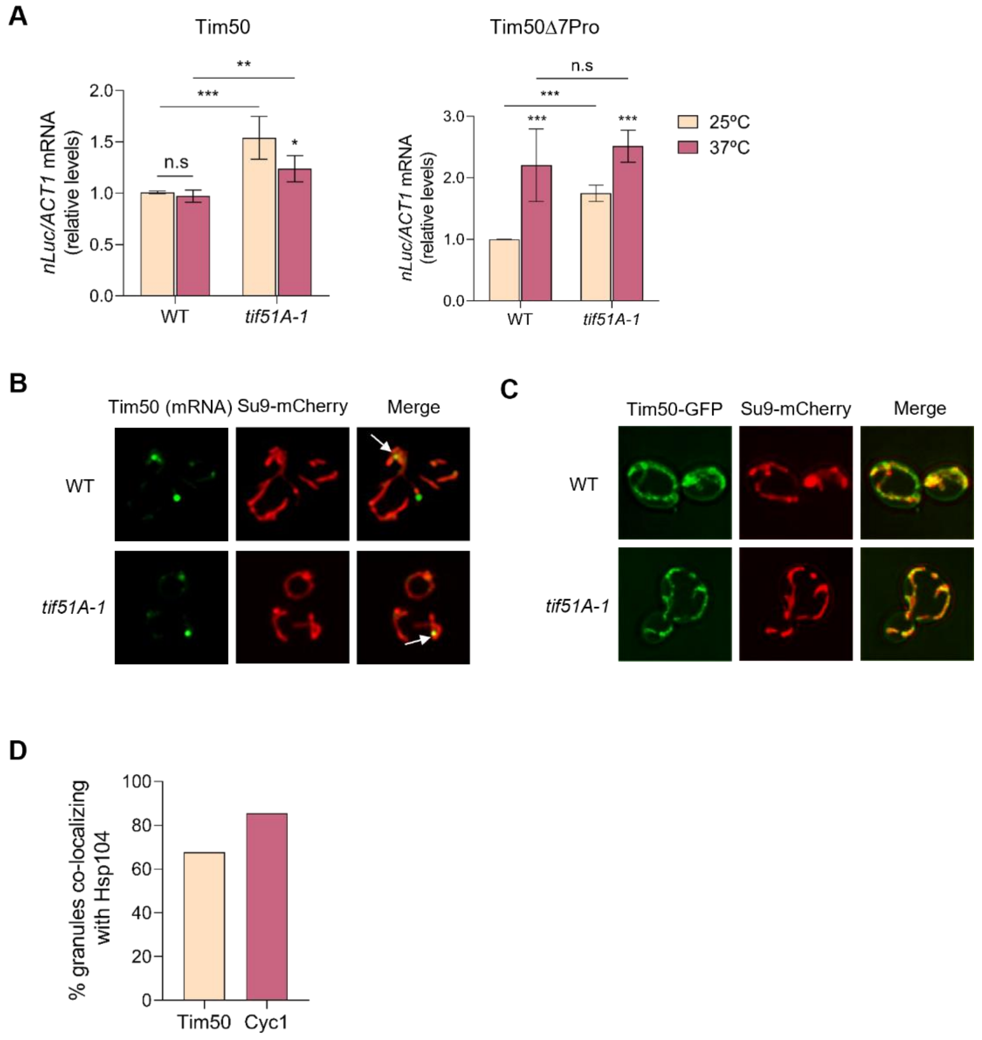


**Figure EV5. eIF5A depletion does not affect *TIM50* mRNA levels nor its mitochondrial mRNA localization.**

(A) Wild-type strain and *tif51A-1* expressing the wild-type Tim50 (left) or Tim50Δ7Pro (right) nLuc constructs were cultured in YPD at 25°C or 37°C for 4 h. One hour after addition of doxycycline to induce nanoluciferase expression, nLuc mRNA relative levels were determined by RT-qPCR.

(B) Wild-type strain and *tif51A-1* were cultured in SGal medium until reaching post-diauxic phase at 25°C, transferred to 37°C for 4 h and then subjected to phase contrast and fluorescence microscopy. Mitochondria were visualized by Su9-mCherry and *TIM50* mRNAs were visualized by the single molecule MS2 tag system.

(C) Wild-type strain and *tif51A-1* expressing Tim50-GFP and Su9-mCherry were cultured in SGal medium at 25°C until reaching post-diauxic phase and subjected to fluorescence microscopy.

(D) Quantification of Tim50 and Cyc1 aggregates co-localizing with Hsp104 from at least 150 cells.

Data information: In (A) Results are presented as mean ± SD from three independent experiments. The statistical significance was measured by using a two-tailed paired Student t-test relative to 25ºC. $^{*}p < 0.05$, $^{**}p < 0.01$, $^{***}p < 0.001$.

**Table S1. Mitochondrial proteins detected in the proteomic analysis with a statistically different 41°C/25°C ratio in at least one eIF5A temperature-sensitive mutant with respect to wild-type.** Proteins are listed according to their functional category. The p-value was determined by a Student's t-test.

| | Gene | Strain | Relative 41/25 protein ratio vs WT [1] | p-value | Location [2] | Description [2] | Putative eIF5A motifs [3] |
|---|---|---|---|---|---|---|---|
| Transport | TIM50 * | *tif51A-1* | 0.824 | 0.2719 | MIM | Mitochondrial import inner membrane translocase | 9 |
| | | *tif51A-3* | 0.862 | 0.2747 | | | |
| | TOM70 | *tif51A-1* | 0.676 | 0.0206 | MOM | Component of the TOM (translocase of outer membrane) complex | 4 |
| | | *tif51A-3* | 0.696 | 0.0299 | | | |
| | MIR1 | *tif51A-1* | 0.660 | 0.0022 | MIM | Mitochondrial phosphate carrier; imports inorganic phosphate into mitochondria | 3 |
| | | *tif51A-3* | 0.508 | 0.0006 | | | |
| | PET9 | *tif51A-1* | 0.605 | 0.0050 | MIM | Major ADP/ATP carrier of the mitochondrial inner membrane | 4 |
| | | *tif51A-3* | 0.479 | 0.0010 | | | |
| | POR1 | *tif51A-1* | 0.551 | 0.0017 | MOM | Mitochondrial porin required for mitochondrial osmotic stability and permeability | 1 |
| | | *tif51A-3* | 0.427 | 0.0013 | | | |
| | TOM20 | *tif51A-1* | 0.279 | 0.0011 | MOM | Component of the TOM (translocase of outer membrane) complex | 2 |
| | | *tif51A-3* | 0.347 | 0.0012 | | | |
| OXPHOS | QCR2 | *tif51A-1* | 0.796 | 0.0480 | MIM | Subunit 2 of ubiquinol cytochrome-c reductase (Complex III) | 1 |
| | | *tif51A-3* | 0.703 | 0.0068 | | | |
| | ATP1 | *tif51A-1* | 0.767 | 0.0014 | MIM | Alpha subunit of the F1 sector of mitochondrial F1F0 ATP synthase | 3 |
| | | *tif51A-3* | 0.692 | 0.0029 | | | |
| | ATP2 | *tif51A-1* | 0.751 | 0.0026 | MIM | Beta subunit of the F1 sector of mitochondrial F1F0 ATP synthase | 5 |
| | | *tif51A-3* | 0.701 | 0.0039 | | | |
| | ATP3 | *tif51A-1* | 0.713 | 0.0451 | MIM | Gamma subunit of the F1 sector of mitochondrial F1F0 ATP synthase | 1 |
| | | *tif51A-3* | 0.758 | 0.0419 | | | |
| | ATP11 | *tif51A-1* | 0.703 | 0.0057 | MIM | Molecular chaperone; required for assembly of mitochondrial F1F0 ATP synthase | 2 |
| | | *tif51A-3* | 0.912 | 0.1657 | | | |
| | COR1 | *tif51A-1* | 0.614 | 0.0005 | MIM | Core subunit of the ubiquinol-cytochrome c reductase complex | 0 |
| | | *tif51A-3* | 0.473 | 0.0001 | | | |
| | CYC1 * | *tif51A-1* | 0.528 | 0.1694 | IMS | Cytochrome c. electron carrier of mitochondrial intermembrane space | 0 |
| | | *tif51A-3* | 0.464 | 0.1309 | | | |
| TCA | ACO1 | *tif51A-1* | 1.029 | 0.7081 | MATRIX | Aconitase; also independently required for mitochondrial genome maintenance | 7 |
| | | *tif51A-3* | 0.825 | 0.0494 | | | |
| | KGD1 | *tif51A-1* | 0.977 | 0.8031 | MATRIX | Subunit of the mitochondrial alpha-ketoglutarate dehydrogenase complex | 2 |
| | | *tif51A-3* | 0.772 | 0.0213 | | | |
| | CIT1 | *tif51A-1* | 0.854 | 0.3034 | MATRIX | Mitochondrial citrate synthase; condenses acetyl-coA and oxaloacetate to form citrate | 1 |
| | | *tif51A-3* | 0.480 | 4.6E-05 | | | |
| | IDH1 | *tif51A-1* | 0.571 | 0.0105 | MATRIX | Subunit of mitochondrial NAD(+)-dependent isocitrate dehydrogenase | 3 |
| | | *tif51A-3* | 0.662 | 0.0241 | | | |
| | LSC2 | *tif51A-1* | 0.534 | 0.0106 | MATRIX | Beta subunit of succinyl-CoA ligase | 4 |
| | | *tif51A-3* | 0.299 | 0.0047 | | | |
| Translation | THS1 | *tif51A-1* | 1.376 | 0.0345 | | Threonyl-tRNA synthetase; essential cytoplasmic protein | 3 |
| | | *tif51A-3* | 1.326 | 0.0325 | | | |
| | BAT1 | *tif51A-1* | 1.314 | 0.0483 | MATRIX | Mitochondrial branched-chain amino acid (BCAA) aminotransferase | 2 |
| | | *tif51A-3* | 1.256 | 0.0410 | | | |
| | TEF4 | *tif51A-1* | 1.273 | 0.0585 | | Gamma subunit of translational elongation factor eEF1B | 1 |
| | | *tif51A-3* | 1.287 | 0.0304 | | | |
| | SNL1 | *tif51A-1* | 1.238 | 0.3920 | | Ribosome-associated protein; proposed to act in protein synthesis | 0 |
| | | *tif51A-3* | 1.341 | 0.0468 | | | |
| | VAS1 | *tif51A-1* | 1.124 | 0.0876 | | Mitochondrial and cytoplasmic valyl-tRNA synthetase | 11 |
| | | *tif51A-3* | 1.227 | 0.0198 | | | |
| | ALA1 | *tif51A-1* | 1.201 | 0.0470 | | Cytoplasmic and mitochondrial alanyl-tRNA synthetase | 9 |
| | | *tif51A-3* | 1.370 | 0.0276 | | | |
| | ILV5 | *tif51A-1* | 1.105 | 0.0649 | MATRIX | Acetohydroxyacid reductoisomerase and mtDNA binding protein | 2 |
| | | *tif51A-3* | 1.267 | 0.0022 | | | |
| | ILV2 | *tif51A-1* | 0.780 | 0.0558 | | Acetolactate synthase; catalyses the first step in isoleucine and valine biosynthesis | 6 |
| | | *tif51A-3* | 0.827 | 0.0362 | | | |

| | Gene | Strain | Relative 41/25 protein ratio vs WT [1] | p-value | Location [2] | Description [2] | Putative eIF5A motifs [3] |
|---|---|---|---|---|---|---|---|
| | TMA19 | *tif51A-1* | 0.677 | 0.0769 | MS | Protein associated with ribosomes; homolog of translationally controlled tumor protein | 1 |
| | | *tif51A-3* | 0.634 | 0.0408 | | | |
| | ILV6 | *tif51A-1* | 0.666 | 0.0185 | MATRIX | Acetolactate synthase. which catalyzes branched-chain amino acid biosynthesis | 6 |
| | | *tif51A-3* | 0.638 | 0.0135 | | | |
| | TUM1 | *tif51A-1* | 0.651 | 0.0370 | | Rhodanese domain sulfur transferase; transfers persulfite from Nfs1p to Uba4p | 0 |
| | | *tif51A-3* | 0.810 | 0.1804 | | | |
| | APE2 | *tif51A-1* | 0.548 | 0.0002 | | Aminopeptidase yscII; may have role in obtaining leucine from dipeptide substrates | 6 |
| | | *tif51A-3* | 0.534 | 0.0003 | | | |
| | HYP2 | *tif51A-1* | 0.284 | 0.0039 | | Translation elongation factor eIF-5A | 1 |
| | | *tif51A-3* | 0.237 | 0.0029 | | | |
| Chaperons | HSP78 | *tif51A-1* | 2.093 | 0.0191 | MATRIX | Oligomeric mitochondrial matrix chaperone; cooperates with Ssc1p after heat shock | 6 |
| | | *tif51A-3* | 1.631 | 0.2857 | | | |
| | HSC82 | *tif51A-1* | 1.342 | 0.0408 | | Cytoplasmic chaperone of the Hsp90 family; plays a role in determining prion variants | 2 |
| | | *tif51A-3* | 1.130 | 0.1624 | | | |
| | HSP10 | *tif51A-1* | 0.884 | 0.8337 | MATRIX | Mitochondrial matrix co-chaperonin; inhibits the ATPase activity of Hsp60p | 0 |
| | | *tif51A-3* | 0.496 | 0.0040 | | | |
| | HSP60 | *tif51A-1* | 0.586 | 0.0475 | MATRIX | Tetradecameric mitochondrial chaperonin | 7 |
| | | *tif51A-3* | 0.503 | 0.0031 | | | |
| | SSC1 | *tif51A-1* | 0.575 | 0.0124 | MIM | A motor component of the translocase of the Inner Mitochondrial membrane | 3 |
| | | *tif51A-3* | 0.676 | 0.0068 | | | |
| Oxidative stress | YHB1 | *tif51A-1* | 1.124 | 0.3646 | MATRIX | Nitric oxide oxidoreductase; plays role in oxidative and nitrosative stress responses | 2 |
| | | *tif51A-3* | 1.527 | 0.0151 | | | |
| | MCR1 | *tif51A-1* | 0.835 | 0.1207 | MOM; IMS | Mitochondrial NADH-cytochrome b5 reductase; involved in ergosterol biosynthesis | 2 |
| | | *tif51A-3* | 0.626 | 0.0032 | | | |
| | SOD2 | *tif51A-1* | 0.824 | 0.3115 | MATRIX | Mitochondrial manganese superoxide dismutase | 3 |
| | | *tif51A-3* | 0.446 | 0.0023 | | | |
| | CCP1 | *tif51A-1* | 0.767 | 0.1371 | IMS | Mitochondrial cytochrome-c peroxidase; degrades reactive oxygen species | 3 |
| | | *tif51A-3* | 0.533 | 0.0147 | | | |
| | CIR2 | *tif51A-1* | 0.668 | 0.0975 | | Putative ortholog of human ETF-dH; may have a role in oxidative stress response | 8 |
| | | *tif51A-3* | 0.482 | 0.0371 | | | |
| | UTH1 | *tif51A-1* | 0.516 | 0.0875 | MIM | Mitochondrial inner membrane protein; role in mitophagy is disputed | 0 |
| | | *tif51A-3* | 0.740 | 0.0438 | | | |
| | HYR1 | *tif51A-1* | 0.444 | 0.0685 | IMS | Glutathione peroxidase; functions as hydroperoxide receptor | 3 |
| | | *tif51A-3* | 0.326 | 0.0391 | | | |
| Others | PDR5 | *tif51A-1* | 2.151 | 0.0006 | | Plasma membrane ATP-binding cassette (ABC) transporter | 3 |
| | | *tif51A-3* | 3.851 | 0.0030 | | | |
| | ECM16 | *tif51A-1* | 2.134 | 0.4299 | | Essential DEAH-box ATP-dependent RNA helicase specific to U3 snoRNP | 9 |
| | | *tif51A-3* | 2.696 | 0.0428 | | | |
| | DBP2 | *tif51A-1* | 1.810 | 0.0731 | | ATP-dependent RNA helicase of the DEAD-box protein family | 4 |
| | | *tif51A-3* | 1.504 | 0.0237 | | | |
| | SNQ2 | *tif51A-1* | 1.665 | 0.0403 | | Plasma membrane ATP-binding cassette (ABC) transporter | 9 |
| | | *tif51A-3* | 1.874 | 0.0070 | | | |
| | MBF1 | *tif51A-1* | 1.559 | 0.1826 | | Transcriptional coactivator | 0 |
| | | *tif51A-3* | 1.493 | 0.0421 | | | |
| | ABF2 | *tif51A-1* | 1.480 | 0.0308 | MATRIX | Mitochondrial DNA-binding protein; involved in DNA replication and recombination | 1 |
| | | *tif51A-3* | 1.403 | 0.0514 | | | |
| | RIB4 | *tif51A-1* | 1.335 | 0.0022 | IMS | Lumazine synthase; catalyzes synthesis of immediate precursor to riboflavin | 1 |
| | | *tif51A-3* | 1.487 | 0.0004 | | | |
| | IDP1 | *tif51A-1* | 1.280 | 0.0194 | MATRIX | Mitochondrial NADP-specific isocitrate dehydrogenase | 2 |
| | | *tif51A-3* | 1.527 | 0.0058 | | | |
| | HXK2 | *tif51A-1* | 1.268 | 0.0934 | MS | Hexokinase isoenzyme 2; phosphorylates glucose in cytosol | 2 |
| | | *tif51A-3* | 1.540 | 0.0198 | | | |
| | LSP1 | *tif51A-1* | 1.185 | 0.0277 | MOM | Eisosome core component | 5 |
| | | *tif51A-3* | 0.837 | 0.0827 | | | |
| | FMN1 | *tif51A-1* | 1.182 | 0.3869 | MIM | Riboflavin kinase. produces riboflavin monophosphate (FMN) | 1 |
| | | *tif51A-3* | 0.634 | 0.0387 | | | |
| | NAT1 | *tif51A-1* | 1.182 | 0.0310 | | Subunit of protein N-terminal acetyltransferase NatA | 3 |
| | | *tif51A-3* | 1.084 | 0.3228 | | | |
| | HEM1 | *tif51A-1* | 1.146 | 0.4233 | MATRIX | 5-aminolevulinate synthase; catalyzes the first step in the heme biosynthetic pathway | 5 |
| | | *tif51A-3* | 0.595 | 0.0305 | | | |

| | Gene | Strain | Relative 41/25 protein ratio vs WT [1] | p-value | Location [2] | Description [2] | Putative eIF5A motifs [3] |
|---|---|---|---|---|---|---|---|
| Others | FPR1 | *tif51A-1* | 1.024 | 0.9786 | | Peptidyl-prolyl cis-trans isomerase; acts as chaperone to prevent protein aggregation. | 5 |
| | | *tif51A-3* | 0.398 | 0.0012 | | | |
| | PGK1 | *tif51A-1* | 1.107 | 0.4701 | | 3-phosphoglycerate kinase; transfers phosphoryl groups to ADP to produce ATP | 6 |
| | | *tif51A-3* | 1.248 | 0.0119 | | | |
| | PRE6 | *tif51A-1* | 0.927 | 0.2263 | MS | Alpha 4 subunit of the 20S proteasome | 4 |
| | | *tif51A-3* | 0.791 | 0.0174 | | | |
| | GPM1 | *tif51A-1* | 0.875 | 0.0540 | IMS | Tetrameric phosphoglycerate mutase; participates in gluconegenesis | 5 |
| | | *tif51A-3* | 0.835 | 0.0253 | | | |
| | NCE102 | *tif51A-1* | 0.866 | 0.2662 | | Protein involved in regulation of pheromone response and mating | 0 |
| | | *tif51A-3* | 0.646 | 0.0199 | | | |
| | OYE2 | *tif51A-1* | 0.853 | 0.1319 | | Conserved NADPH oxidoreductase containing flavin mononucleotide (FMN) | 3 |
| | | *tif51A-3* | 0.754 | 0.0151 | | | |
| | DUG1 | *tif51A-1* | 0.839 | 0.0057 | | Cys-Gly metallo-di-peptidase | 6 |
| | | *tif51A-3* | 0.799 | 0.0592 | | | |
| | FBA1 | *tif51A-1* | 0.839 | 0.0181 | MS | Fructose 1.6-bisphosphate aldolase; required for glycolysis and gluconeogenesis | 2 |
| | | *tif51A-3* | 0.984 | 0.6094 | | | |
| | DNM1 | *tif51A-1* | 0.830 | 0.0329 | MOM | Dynamin-related GTPase involved in mitochondrial organization | 8 |
| | | *tif51A-3* | 0.849 | 0.0464 | | | |
| | YIM1 | *tif51A-1* | 0.811 | 0.2762 | | Aldehyde reductase; involved in detoxification of lignocellulose-derived aldehydes | 5 |
| | | *tif51A-3* | 0.474 | 0.0294 | | | |
| | YDL086W | *tif51A-1* | 0.810 | 0.1067 | | Putative carboxymethylenebutenolidase | 0 |
| | | *tif51A-3* | 0.729 | 0.0442 | | | |
| | PIL1 | *tif51A-1* | 0.812 | 0.0610 | MOM | Eisosome core component involved in endocytosis | 3 |
| | | *tif51A-3* | 0.694 | 0.0230 | | | |
| | FRD1 | *tif51A-1* | 0.802 | 0.0436 | | Soluble fumarate reductase; may interact with ribosomes | 3 |
| | | *tif51A-3* | 0.898 | 0.2154 | | | |
| | GNP1 | *tif51A-1* | 0.797 | 0.3180 | | Broad specificity amino acid permease; major serine permease | 5 |
| | | *tif51A-3* | 0.534 | 0.0442 | | | |
| | LAT1 | *tif51A-1* | 0.795 | 0.0100 | MATRIX | E2 component of the pyruvate dehydrogenase complex | 0 |
| | | *tif51A-3* | 0.683 | 0.0092 | | | |
| | MSS116 | *tif51A-1* | 0.787 | 0.0269 | MATRIX | Mitochondrial transcription elongation factor; DEAD-box protein | 1 |
| | | *tif51A-3* | 1.019 | 0.7656 | | | |
| | ARG7 | *tif51A-1* | 0.785 | 0.0507 | MATRIX | Mitochondrial ornithine acetyltransferase; catalyzes the fifth step in arginine biosynthesis | 0 |
| | | *tif51A-3* | 0.771 | 0.0406 | | | |
| | SEC4 | *tif51A-1* | 0.775 | 0.0250 | MOM | Rab family GTPase; essential for vesicle-mediated exocytic secretion and autophagy | 0 |
| | | *tif51A-3* | 0.694 | 0.0158 | | | |
| | DPM1 | *tif51A-1* | 0.769 | 0.0576 | MOM | Dolichol phosphate mannose synthase of ER membrane | 0 |
| | | *tif51A-3* | 0.759 | 0.0249 | | | |
| | VPS21 | *tif51A-1* | 0.769 | 0.0279 | MOM | Endosomal Rab family GTPase | 0 |
| | | *tif51A-3* | 0.669 | 0.0055 | | | |
| | CYS4 | *tif51A-1* | 0.748 | 0.0885 | | Cystathionine beta-synthase; catalyzes the first committed step in cysteine biosynthesis | 1 |
| | | *tif51A-3* | 0.726 | 0.0253 | | | |
| | NUP2 | *tif51A-1* | 0.747 | 0.0102 | | Nucleoporin involved in nucleocytoplasmic transport | 5 |
| | | *tif51A-3* | 0.678 | 0.0553 | | | |
| | YKT6 | *tif51A-1* | 0.747 | 0.0455 | | Vesicle membrane protein (v-SNARE) with acyltransferase activity | 0 |
| | | *tif51A-3* | 0.739 | 0.0499 | | | |
| | PMA1 | *tif51A-1* | 0.739 | 0.0202 | MIM | Plasma membrane P2-type H+-ATPase; pumps prótons out of the cell | 4 |
| | | *tif51A-3* | 0.713 | 0.0167 | | | |
| | ECM33 | *tif51A-1* | 0.738 | 0.0055 | | GPI-anchored protein involved in efficient glucose uptake | 3 |
| | | *tif51A-3* | 0.660 | 0.0165 | | | |
| | RAD23 | *tif51A-1* | 0.718 | 0.0520 | | Proteasome-associated ubiquitin receptor; recruits substrates to the proteasome | 4 |
| | | *tif51A-3* | 0.541 | 0.0123 | | | |
| | YCP4 | *tif51A-1* | 0.717 | 0.0360 | | Protein of unknown function; has sequence and structural similarity to flavodoxins | 3 |
| | | *tif51A-3* | 0.709 | 0.0290 | | | |
| | CPR3 | *tif51A-1* | 0.719 | 0.2164 | | Mitochondrial peptidyl-prolyl cis-trans isomerasematr | 0 |
| | | *tif51A-3* | 0.504 | 0.0106 | | | |
| | PRX1 | *tif51A-1* | 0.709 | 0.0189 | MATRIX | Mitochondrial peroxiredoxin with thioredoxin peroxidase activity | 2 |
| | | *tif51A-3* | 0.572 | 0.0028 | | | |

| | Gene | Strain | Relative 41/25 protein ratio vs WT [1] | p-value | Location [2] | Description [2] | Putative eIF5A motifs [3] |
|---|---|---|---|---|---|---|---|
| Others | VPS1 | *tif51A-1* | 0.680 | 0.0009 | MOM | Dynamin-like GTPase required for vacuolar sorting | 6 |
| | | *tif51A-3* | 0.719 | 0.0077 | | | |
| | ENO1 | *tif51A-1* | 0.672 | 0.0372 | | Enolase I. a phosphopyruvate hydratase | 2 |
| | | *tif51A-3* | 0.666 | 0.0293 | | | |
| | YPT7 | *tif51A-1* | 0.669 | 0.0362 | | Enolase I. a phosphopyruvate hydratase | 0 |
| | | *tif51A-3* | 1.020 | 0.8286 | | | |
| | AIM45 | *tif51A-1* | 0.667 | 0.0219 | | Putative ortholog of mammalian ETF-alpha; interacts with frataxin | 4 |
| | | *tif51A-3* | 0.908 | 0.7639 | | | |
| | FAA1 | *tif51A-1* | 0.589 | 0.0120 | MOM | Long chain fatty acyl-CoA synthetase; activates fatty acids | 10 |
| | | *tif51A-3* | 0.482 | 0.0046 | | | |
| | RTN1 | *tif51A-1* | 0.579 | 0.0008 | | Reticulon protein; involved in nuclear pore assembly and tubular ER morphology | 0 |
| | | *tif51A-3* | 0.598 | 0.0007 | | | |
| | LAP3 | *tif51A-1* | 0.573 | 0.0011 | | Cysteine aminopeptidase with homocysteine-thiolactonase activity | 2 |
| | | *tif51A-3* | 0.606 | 0.0108 | | | |
| | DCS1 | *tif51A-1* | 0.568 | 0.0137 | | Non-essential hydrolase involved in mRNA decapping; activates Xrn1p | 1 |
| | | *tif51A-3* | 0.428 | 0.0057 | | | |
| | YME1 | *tif51A-1* | 0.509 | 0.0266 | MIM | Catalytic subunit of i-AAA protease complex; helps degradation of unfolded gene products. | 7 |
| | | *tif51A-3* | 0.682 | 0.1948 | | | |
| | ADK1 | *tif51A-1* | 0.505 | 0.0859 | IMS | Adenylate kinase. required for purine metabolism; controls ATP homeostasis | 3 |
| | | *tif51A-3* | 0.359 | 0.0281 | | | |
| | DOP1 | *tif51A-1* | 0.502 | 0.0308 | | Protein involved in vesicular transport at trans-Golgi network | 4 |
| | | *tif51A-3* | 0.391 | 0.0120 | | | |
| | SOD1 | *tif51A-1* | 0.502 | 0.1241 | IMS | Cytosolic copper-zinc superoxide dismutase and sulfide oxidase | 0 |
| | | *tif51A-3* | 0.341 | 0.0373 | | | |
| | EIS1 | *tif51A-1* | 0.486 | 0.0432 | | Component of the eisosome required for proper eisosome assembly | 4 |
| | | *tif51A-3* | 0.415 | 0.0453 | | | |
| | CFT1 | *tif51A-1* | 0.478 | 0.0407 | | RNA-binding subunit of the mRNA cleavage and polyadenylation factor | 8 |
| | | *tif51A-3* | 0.706 | 0.2263 | | | |
| | PST2 | *tif51A-1* | 0.470 | 0.0042 | | FMN-dependent NAD(P)H: quinone oxidoreductase | 4 |
| | | *tif51A-3* | 0.366 | 0.0011 | | | |
| | ZEO1 | *tif51A-1* | 0.370 | 0.1195 | MOM | Peripheral membrane protein of the plasma membrane; regulates cell integrity | 0 |
| | | *tif51A-3* | 0.294 | 0.0473 | | | |
| | ALD4 | *tif51A-1* | 0.333 | 0.0015 | MATRIX | Mitochondrial aldehyde dehydrogenase | 3 |
| | | *tif51A-3* | 0.182 | 0.0005 | | | |
| | HXT7 | *tif51A-1* | 0.329 | 0.0266 | | FMN-dependent NAD(P)H:quinone oxidoreductase; induced by oxidative stress | 2 |
| | | *tif51A-3* | 0.281 | 0.0244 | | | |
| | TDH1 | *tif51A-1* | 0.310 | 0.0002 | | Glyceraldehyde-3-phosphate dehydrogenase; involved in glycolysis and gluconeogenesis | 2 |
| | | *tif51A-3* | 0.377 | 4E-04 | | | |

[1] Relative 41°C/25°C protein ratio of the indicated eIF5A mutant strain respect to the wild-type.

[2] Information obtained from SGD (Saccharomyces Genome Database). MS, mitochondrial surface; MOM, mitochondrial outer membrane; MIM, mitochondrial inner membrane; IMS, mitochondrial intermembrane space.

[3] Number of eIF5A-dependent tripeptide motifs described in Pelechano and Alepuz, 2017.

* Down-regulated proteins in eIF5A mutants respect to wild-type but with a non-significant statistical value.

**Table S2. Yeast strains used in this study**

| Name | Genotype | Source |
|---|---|---|
| **BY4741** | MATa *ura3Δ0 leu2Δ0 his3Δ1 met15Δ0* | Euroscarf |
| ***tif51A-1*** | BY4741 MATa *ura3Δ0 leu2Δ0 his3Δ1 met15Δ0 tif51A-1::kanR* | (Li *et al.,* 2011) |
| ***tif51A-3*** | BY4741 MATa *ura3Δ0 leu2Δ0 his3Δ1 met15Δ0 tif51A-3::kanR* | (Li *et al.,* 2011) |
| **PAY864** | BY4741 MATa *ura3Δ0 leu2Δ0 his3Δ1 met15Δ0 TIM50-3HA-his3MX6* | This study |
| **PAY866** | BY4741 MATa *ura3Δ0 leu2Δ0 his3Δ1 met15Δ0 tif51A-1::kanR TIM50-3HA-his3MX6* | This study |
| **PAY937** | BY4741 MATa *ura3Δ0 leu2Δ0 his3Δ1 met15Δ0 PDR5-GFP-his3MX6* | This study |
| **PAY938** | BY4741 MATa *ura3Δ0 leu2Δ0 his3Δ1 met15Δ0 tif51A-1::kanR PDR5-GFP-his3MX6* | This study |
| **PAY1066** | BY4741 MATa *ura3Δ0 leu2Δ0 his3Δ1 met15Δ0 TOM70-GFP-his3MX6* | This study |
| **PAY1067** | BY4741 MATa *ura3Δ0 leu2Δ0 his3Δ1 met15Δ0 tif51A-1::kanR TOM70-GFP-his3MX6* | This study |
| **PAY1068** | BY4741 MATa *ura3Δ0 leu2Δ0 his3Δ1 met15Δ0 CYC1-GFP-his3MX6* | This study |
| **PAY1069** | BY4741 MATa *ura3Δ0 leu2Δ0 his3Δ1 met15Δ0 tif51A-1::kanR CYC1-GFP-his3MX6* | This study |
| **PAY1072** | BY4741 MATa *ura3Δ0 leu2Δ0 his3Δ1 met15Δ0 ILV2-GFP-his3MX6* | This study |
| **PAY1073** | BY4741 MATa *ura3Δ0 leu2Δ0 his3Δ1 met15Δ0 tif51A-1::kanR ILV2-GFP-his3MX6* | This study |
| **PAY1078** | BY4741 MATa *ura3Δ0 leu2Δ0 his3Δ1 met15Δ0 TIM50-GFP-his3MX6* | This study |
| **PAY1079** | BY4741 MATa *ura3Δ0 leu2Δ0 his3Δ1 met15Δ0 tif51A-1::kanR TIM50-GFP-his3MX6* | This study |
| **PAY1080** | BY4741 MATa *ura3Δ0 leu2Δ0 his3Δ1 met15Δ0 YTA12-GFP-his3MX6* | This study |
| **PAY1081** | BY4741 MATa *ura3Δ0 leu2Δ0 his3Δ1 met15Δ0 tif51A-1::kanR YTA12-GFP-his3MX6* | This study |
| **PAY1085** | BY4741 MATa *ura3Δ0 leu2Δ0 his3Δ1 met15Δ0 TIM50Δ7Pro-GFP-his3MX6* | This study |
| **PAY1086** | BY4741 MATa *ura3Δ0 leu2Δ0 his3Δ1 met15Δ0 tif51A-1::kanR TIM50Δ7Pro-GFP-his3MX6* | This study |
| **PAY1107** | BY4741 MATa *ura3Δ0 leu2Δ0 his3Δ1 met15Δ0 TIM50-GFP-his3MX6 HSP104-RFP-clonNAT* | This study |
| **PAY1113** | BY4741 MATa *ura3Δ0 leu2Δ0 his3Δ1 met15Δ0 tif51A-1::kanR TIM50-GFP-his3MX6 HSP104-RFP-clonNAT* | This study |
| **PAY1142** | BY4741 MATa *ura3Δ0 leu2Δ0 his3Δ1 met15Δ0 CYC1-GFP-his3MX6 HSP104-RFP-clonNAT* | This study |
| **PAY1144** | BY4741 MATa *ura3Δ0 leu2Δ0 his3Δ1 met15Δ0 tif51A-1::kanR CYC1-GFP-his3MX6 HSP104-RFP-clonNAT* | This study |

**Table S3. Plasmids used in this study**

| Name | Plasmid description | Source |
|---|---|---|
| **PA201** | pFA6a-3HA-HIS3MX6 | (Longtine *et al.,* 1998) |
| **PA242** | pFA6a-GFP-HIS3MX6 | (Longtine *et al.,* 1998) |
| **PA354** | pMK46-IAA17-kanMX | Dr. Ethel Queralt |
| **PA367** | pYM43-Redstar2-clonNAT | Euroscarf |
| **PA386** | pYES2-pGAL-FLAG-htt25QP-GFP-URA3 | (Berglund *et al.*, 2017) |
| **PA388** | pYES2-pGAL-FLAG-TIM50-GFP-URA3 | This study |
| **TTP76** | pRS406-GPDp-Su9-mCherry-URA3 | Dr. Brian M. Zid |
| **TTP80** | pRS405-CYC1p-MS2-4xGFP-LEU2 | Dr. Brian M. Zid |
| **TTP145** | pRS403 TIM50p-TIM50mts(1-300)-TIM50orf-flagiRFP-TIM50ter-MS2tag | Dr. Brian M. Zid |
| **ZP447** | pAG306-ptetO$_7$-TIM505 'UTR-TIM50-nLuc-CYC1term-URA3 | Dr. Brian M. Zid |
| **ZP448** | pAG306-ptetO$_7$-TIM505 'UTR-TIM50Δ7Pro-nLuc-CYC1term-URA3 | Dr. Brian M. Zid |
| **ZP562** | pAG306-ptetO$_7$-TIM505 'UTR-SDH2-nLuc-CYC1term-URA3 | Dr. Brian M. Zid |
| **ZP603** | pAG306-ptetO$_7$-TIM505 'UTR-CYC1-nLuc-CYC1term-URA3 | This study |
| **ZP605** | pAG306-ptetO$_7$-TIM505 'UTR-COX5A-nLuc-CYC1term-URA3 | This study |

**Table S4. Oligonucleotides used in this study**

| Primer | Sequence (5'-3') |
|---|---|
| **Gene expression detection by RT-qPCR** | |
| ACT1-F | TCGTTCCAATTTACGCTGGTT |
| ACT1-R | CGGCCAAATCGATTCTCAA |
| ATP1-F | AGACCTGCCATTAACGTTGG |
| ATP1-R | AGCAAAAGCAGCGACTTCTC |
| CIS1-F | TGCAGAGTGGGTAGCATGTC |
| CIS1-R | TGGGCAGCCTTGAGTAAATC |
| COX5A-F | ATCCAGATGGGAGAACATGC |
| COX5A-R | AGCTTGCTTTTCAGGCTCAG |
| CYC1-F | AGATGTCTACAATGCCACACC |
| CYC1-R | CCCTTCAGCTTGACCAGAGT |
| EFT2-F | TGTTCATCAAGGCCATTCAA |
| EFT2-R | GTTACCGGCTGGACAGTCAT |
| GFP-F | CACATGAAGCAGCACGACTT |
| GFP-R | GGTCTTGTAGTTGCCGTCGT |
| GRE2-F | GCCTTCCAAAAGAGGGAAAC |
| GRE2-R | ATGGGTAGCACCAGAACCTG |
| HSP60-F | CACTGATCCAAAGTCGAGCA |
| HSP60-R | CAAGAGCTTCACCGTCAACA |
| MSP1-F | ACGGCAACCTTAAAAGCTGA |
| MSP1-R | CCTCCTCAAAAAACGCATCAT |
| NLUC-F | AGAACAAGGTGGTGTTTCTTCT |
| NLUC-R | CCCATTTGATCACCAGATAAACCT |
| PDR1-F | TCCAAATGCGAGATTTTTCC |
| PDR1-R | CGAAGATGGGGTTGAAGGTA |
| PDR3-F | AGATGGGATTGTCTCGTTGG |
| PDR3-R | CTGAAATCCTTCGGCAAGAG |
| PDR5-F | GTACCGTGGTTGGAGCTGTT |
| PDR5-R | GAAACCACGCCATTTGTCTT |
| PDR15-F | CCAAGGTCGGAAACGATCTA |
| PDR15-R | CAATGTCAGCCTGGGTTTTT |
| PET9-F | AGGCCATGTTTGGTTTCAAG |
| PET9-R | TCAAACCGTTGAATTGACGA |
| POR1-F | AAACCGGCTTGGGTCTAACT |
| POR1-R | GACGCCTGGAGTCAAAGAAG |
| RPP2B-F | GGAAGGTAAGGGCTCTTTGG |
| RPP2B-R | TTCTTCAGCAGCATCACCAC |
| TIM50-F | TCTGCGTTGACAGGTACTGC |
| TIM50-R | AATCAGGGAAAGGTGGCTCT |

| | |
|---|---|
| TOM20-F | CCGCAATTCAGGAAAGTGTT |
| TOM20-R | CCTTTTGCAGCTTCACTTCC |
| TOM70-F | GACCCAAGAAGTGAGCAAGC |
| TOM70-R | AGCGGCTTCAGCAAAAGTAA |
| **Gene tagging by PCR** | |
| CYC1-F2 | AAAGACAGAAACGACTTAATTACCTACTTGAAAAAAGCCTGTGAGCGGATCCCCGGGTTAATTAA |
| CYC1-R1 | TGACATAACTAATTACATGATATCGACAAAGGAAAAGGGGCCTGTGAATTCGAGCTCGTTTAAAC |
| HSP104-RFP-F | GATGACGATAATGAGGACAGTATGGAAATTGATGATGACCTAGATCGTACGCTGCAGGTCGAC |
| HSP104-RFP-R | TACTGCTTCTTGTTCGAAAGTTTTTTAAAAATCACACTATATTAAATCAATCGATGAATTCGAGCTCG |
| ILV2-F2 | AGACAACAGACTGAATTACGTCATAAGCGTACAGGCGGTAAGCACCGGATCCCCGGGTTAATTAA |
| ILV2-R1 | TGCATTTTTTACTGAAAATGCTTTTGAAATAAATGTTTTTGAAATGAATTCGAGCTCGTTTAAAC |
| PDR5-F2 | TGGTTAGCAAGAGTGCCTAAAAAGAACGGTAAACTCTCCAAGAAACGGATCCCCGGGTTAATTAA |
| PDR5-R1 | GTCCATCTTGGTAAGTTTCTTTTCTTAACCAAATTCAAAATTCTAGAATTCGAGCTCGTTTAAAC |
| TIM50-F2 | TTATTTGAAGAGGAAAAGAAAAAAGAAGAAGATTGCTGAATCCAAACGGATCCCCGGGTTAATTAA |
| TIM50-R1 | CACACATAGATACGTAGATACATGAGAAGAGGGTTTACATGAAAAGAATTCGAGCTCGTTTAAAC |
| TOM70-F2 | AAGATTCAAGAAACTTTAGCTAAATTACGCGAACAGGGTTTAATGCGGATCCCCGGGTTAATTAA |
| TOM70-R1 | TAGTTTTTGTCTTCTCCTAAAAGTTTTTAAGTTTATGTTTACTGTGAATTCGAGCTCGTTTAAAC |
| YTA12-F2 | GAAGAAAAAAACGAAAAACGTAATGAGCCTAAGCCATCTACAAACCGGATCCCCGGGTTAATTAA |
| YTA12-R1 | ATATGTAGAACAGTCTTCCTCCATTTCTTTGTATTGTGAAATATCGAATTCGAGCTCGTTTAAAC |
| **Proline deletion by PCR** | |
| TIM50-delPro-F | CCTACTTCCAAGAGCCACCTTTCCCTGATTTACTACCAAAGGCCATTAACTCTTG |
| TIM50-delPro-R | CACACATAGATACGTAGATACATGAGAAGAGGGTTTACATGAAAA |
| **Cloning into pYES2 plasmid** | |
| TIM50-F3 | AAGCTTGGTACCGCCATGGACTACAAGGACGACGATGACAAGCTGCTGTCCATTTTAAGAAATTC |
| TIM50-R3 | CACCCCGGTGAACAGCTCCTCGCCCTTGCTCACCAGGGATCCCCCTTTGGATTCAGCAATCTTCT |
| **Cloning into ZP446 plasmid** | |
| ZP446-F | ATGGTTTTTACTTTAGAAGATTTTG |
| ZP446-R | TGCAAGCGGGTGATTTTTGGAAGTTTATTCTAGC |
| CYC1-F | CCAAAAATCACCCGCTTGCAATGACTGAATTCAAGGCCGGTTCTGCTAAG |
| CYC1-R | CAACAAAATCTTCTAAAGTAAAAACCATCTCACAGGCTTTTTTCAAGTAGGTAAT |
| COX5A-F | CCAAAAATCACCCGCTTGCAATGTTACGTAACACTTTTACTAGAGCTGGT |
| COX5A-R | CAACAAAATCTTCTAAAGTAAAAACCATTTTAGATTGGACCTGAGAATAACCAC |
